# Therapeutic Potential of Hypoxia-Preconditioned hiPSC-Epicardial Cell-Derived Exosomes in Mice with Myocardial Infarction

**DOI:** 10.64898/2026.04.19.719232

**Authors:** Zhixuan Qiu, Yun Jiang, Pengfei Zhang, Hao Li, Yaoxian Yu, Yanshan Gong, Ling Gao

## Abstract

**Background:** It has been demonstrated that stem cell transplantation promotes healing of the infarcted heart through paracrine effects. However, the therapeutic potential of exosomes secreted by hiPSC-derived epicardial cells (hEP-Exos) for treating infarcted hearts remains unclear. Myocardial infarction (MI) can trigger EP activation, increasing EP paracrine function. Therefore, this study aims to determine and compare the cardioprotective effects of exosomes secreted by hEPs under normoxic (Exo-N) and hypoxic (Exo-H) conditions in MI mice and to explore the underlying mechanisms.

**Methods:** Two types of exosomes were collected by ultracentrifugation and delivered via intramyocardial injection in a murine MI model. The protective effects of Exo-N and Exo-H on the infarcted heart were assessed using echocardiography, histological examination, and immunofluorescence analysis. Additionally, microRNA sequencing, luciferase activity assays, and miRNA gain-and loss-of-function experiments were performed to identify enriched miRNAs and investigate their roles in different exosome populations.

**Results:** In vitro, both Exo-N and Exo-H enhanced the migration and tube-formation capacities in human umbilical vein endothelial cells (HUVECs) and reduced the apoptosis in hiPSC-derived cardiomyocytes (hCMs) under oxygen-glucose deprivation (OGD), with Exo-H exhibiting a stronger effect. In vivo, both Exo-N and Exo-H significantly improved contractile function, reduced infarct size, and mitigated adverse remodeling in mouse hearts with MI, accompanied by increased cardiomyocyte survival and angiogenesis, with Exo-H showing superior efficacy. Mechanistically, miRNA sequencing revealed distinct cargo profiles between Exo-N and Exo-H. miR-214-3p was identified as a key mediator of the enhanced therapeutic potency of Exo-H. miR-214-3p promoted EC angiogenesis by suppressing vasohibin-1 and attenuated cardiomyocyte mitochondrial fission and apoptosis by suppressing mitochondrial elongation factor 2 (MIEF2).

**Conclusions:** This study demonstrates that administration of hEP-Exos, particularly Exo-H, provides robust cardioprotection by enhancing cardiomyocyte survival and angiogenesis, potentially mediated by miR-214-3p. These findings suggest that conditioned hEP-Exos could be a promising and effective acellular therapeutic option for treating MI.

## Introduction

Myocardial infarction (MI) is a critical condition characterized by a significant reduction or complete cessation of blood flow through the coronary arteries, ultimately causing cardiomyocyte death. It is a leading cause of morbidity and mortality worldwide, highlighting the urgent need for new therapeutic strategies to promote myocardial repair ^[1, 2]^. Epicardial cells (EPs), the outermost layer of the heart, play a vital role in heart development and in repair after injury ^[3, 4]^. Studies have shown that MI triggers an epithelial-mesenchymal transition (EMT) in the endogenous epicardium, leading to the activation of EPs and their differentiation into various cardiac lineages ^[5]^. These cells participate in the repair process by replacing lost cells and repairing damaged myocardium through the release of paracrine factors ^[4–6]^. However, EPs constitute only a small fraction of the overall cardiac cell population. To overcome this limitation, studies have shown that transplanting human embryonic stem cell (hESC)-derived EPs into mouse hearts yields significant therapeutic benefits ^[7]^. Most current research has focused on quiescent EPs, whereas the therapeutic potential of activated EPs has received less attention.

Hypoxia can partially mimic the ischemic microenvironment following MI. Previous studies have shown that hypoxic conditions can activate primary epicardial cells ^[8, 9]^, but it is unclear whether this effect also applies to human induced pluripotent stem cell (hiPSC)-derived EPs (hEPs). Furthermore, evidence suggests that hypoxia may enhance paracrine function through mechanisms involving HIF-1α activation and alterations in cargo composition ^[10, 11]^. Extracellular vesicles, including exosomes, have attracted increasing attention for their vital role in mediating the cardioprotective effects of stem cells and their derivatives. Exosomes encapsulate various biomolecules, including proteins, lipids, and nucleic acids (such as mRNA and miRNA), and play a key role in myocardial repair after injury ^[7, 12–14]^. However, the therapeutic potential of exosomes secreted by hEPs (hEP-Exos), especially those activated under hypoxic conditions, for treating infarcted hearts remains unclear and needs to be elucidated.

Mitochondria, often referred to as the “powerhouses” of the cell, are essential organelles responsible for producing energy ^[15]^. Beyond their crucial metabolic functions, mitochondria are highly dynamic organelles that constantly undergo fusion and fission events to regulate their morphology, number, and size ^[16]^. Disruptions in mitochondrial dynamics, caused by factors such as oxidative stress or ischemia, can lead to apoptotic cell death and the opening of the mitochondrial permeability transition pore (mPTP), thereby further exacerbating myocardial injury ^[17]^. Therefore, improving mitochondrial dynamics to restore cellular function may be a key strategy for enhancing cardioprotection after MI. However, the impact of hEP-Exos on this process remains unclear.

In this study, we collected exosomes derived from hEPs cultured under normoxic (Exo-N) or hypoxic (Exo-H) conditions. Through a combination of in vivo and in vitro assessments, we aimed to investigate (i) whether the two types of hEP-Exos can promote angiogenesis and reduce cardiomyocyte apoptosis, thereby enhancing myocardial recovery in mice with MI; (ii) the differences in effectiveness between these two types of exosomes; and (iii) the underlying mechanisms and contributions to infarcted heart healing. This research will support the view that conditioned hEP-Exos may be a beneficial acellular therapeutic option for treating MI.

## Materials and methods

### hiPSC culture, hEP induction, and hypoxia treatment

hiPSCs used in this study were obtained from the GMP Experimental Center of Shanghai East Hospital and the National Stem Cell Transformation Resource Bank. They were reprogrammed from human umbilical cord blood mononuclear cells by transduction with OCT4, SOX2, KLF4, and cMYC. hiPSCs were routinely maintained in mTeSR1 medium (Stem Cell Technologies, #85850, Canada) on Matrigel-coated plates (Corning, 3516, USA) and differentiated into hEPs as described previously ^[18, 19]^. Briefly, hiPSCs were plated onto Matrigel-coated plates at 25,000 cells/cm2 in mTeSR supplemented with 10 µM Y-27632 (StemCell Technologies, #72307). When hiPSCs reached 85% confluence, 6 µM CHIR99021 (StemCell Technologies, #72054) was added to the chemically defined medium (CDM), which contained RPMI-1640 (Thermo Fisher Scientific, 11875119, USA), 213 µg/mL L-ascorbic acid 2-phosphate (Sigma–Aldrich, A8960, Germany), and 2 mg/mL bovine serum albumin (Sigma–Aldrich, A4628) to initiate differentiation. On day 2, the medium was replaced with CDM supplemented with 5 µM IWR-1 (Merck, 1127442-82-3, Germany). On day 4, differentiating cells were dissociated with Accutase (Stem Cell Technologies, 07922) for 5 min, then plated at a 1:24 density in LasR medium (advanced DMEM/F12, Thermo Fisher Scientific, 12634028), supplemented with 1× GlutaMax (Thermo Fisher Scientific, 35050061) and 100 µg/mL ascorbic acid (Sigma–Aldrich, 50-81-7), with 10 µM Y-27632. 24 h later, the cells were incubated with 3 µM CHIR99021 in LasR medium for 2 days; afterward, the medium was changed every 2 days until passaging on day 12. The hEPs were then dissociated with 0.05% Trypsin-EDTA (Thermo Fisher Scientific, 17892) for passaging, cryopreservation, and analysis.

Hypoxic conditioning was performed by placing the cells in an oxygen-controlled cabinet within an incubator maintained at 1% O_2_, 5% CO_2_, and balanced N_2_ in basal medium. Normoxic conditioning of hEPs was performed at 21% O_2_ and 5% CO_2_ in basal medium. Both normoxic and hypoxic conditioned media were collected for exosome isolation.

### Exosome isolation and identification

The exosomes were isolated separately from the normoxic and hypoxic conditioned media, as previously described ^[12, 13, 20]^. In brief, the collected conditioned medium was centrifuged at 300 g for 30 min, then at 2,000 g for 30 min, both at 4 °C to remove cells and dead cells. It was then centrifuged at 10,000 g for 30 min at 4 °C to remove cell debris, and finally centrifuged twice at 120,000 g for 120 min at 4 °C using an Optima+XPN-100 (Beckman). The final pellet containing exosomes was resuspended in PBS. Meanwhile, miR-214-3p antagomir or an equal dose of antagomir negative control (NC) was transfected into hEPs using a Ribo FECT^TM^ CP Transfection Kit (RiboBio, C10511-05, China), followed by hypoxic treatment to collect hypoxic conditioned medium. Exosomes secreted by the transfected hEPs were isolated and identified as Exo-H^anti-miR-214-3p^ or Exo-H^NC^.

The particle-size distributions and yield of exosomes reconstituted in PBS were determined by nanoparticle tracking analysis (NTA) using a NanoSight instrument. The protein content was measured using the MicroBCA Protein Assay Kit (Beyotime, P0012, China), and the expression levels of marker proteins (ALG-2-interacting protein X (Alix), tumor susceptibility gene 101 protein (TSG101), CD63, and CD9) were evaluated by western blot. The ultrastructure of exosomes was assessed by transmission electron microscopy (TEM) as described previously.^[21]^

### Generation of hiPSC-derived cardiomyocytes (hCMs)

hCMs were generated from hiPSCs. In brief, hiPSCs were expanded on Matrigel-coated dishes for 4 days, then treated with 6 μM CHIR99021 (StemCell, Canada) in RPMI 1640 with 2% B27 without insulin for 48 h. Subsequently, cells were exposed to 5 μM IWR-1 in RPMI 1640 with 2% B27 without insulin for 48 h, followed by switching to RPMI 1640 with 2% B27 complete, with medium changed every 3 days. Beating cells typically emerged around 8 days post-differentiation initiation. Purification of hCMs involved culturing in a glucose-depleted medium with ample lactate for 3 days.

### Exosome uptake assessment

To monitor exosome internalization, purified exosomes were labeled with a Dil red fluorescent cell linker kit (Beyotime, C1036) for in vitro studies, following the manufacturer’s instructions. The prelabeled exosomes were then added to hCMs and human umbilical vein endothelial cells (HUVECs), which were subsequently fixed and stained for immunofluorescence.

### Western blot analysis

The cells and exosomes were lysed using radioimmunoprecipitation assay lysis buffer. The proteins were separated by gel electrophoresis, transferred to a polyvinylidene fluoride membrane, and blocked with 5% bovine serum albumin for 2 h. The membranes were incubated overnight with primary antibodies against Alix (Cell Signaling Technology, #2171, USA), TSG101 (Abcam, ab125011, UK), CD63 (Santa Cruz Biotechnology, sc-5275, USA), CD9 (Thermo Fisher Scientific, MA5-31980, USA), vasohibin-1 (VASH1, Proteintech, 12730-1-AP, China), vascular endothelial growth factor A (VEGFA, Proteintech, 26381-1-AP), dynamin-related protein 1 (Drp1, Cell Signaling Technology, # 8570T), voltage-dependent anion channel (VDAC, Proteintech, 55259-1-AP), mitochondrial elongation factor 2 (MIEF2, Proteintech, 26038-1-AP), and glyceraldehyde 3-phosphate dehydrogenase (GAPDH, Proteintech, 60004-1-Ig). The membranes were then incubated with horseradish peroxidase-conjugated secondary antibodies at room temperature for 2 h and exposed via enhanced chemiluminescence.

### Luciferase reporter assay

The VASH1 and MIEF2 sequences were amplified and cloned into the pmirGLO dual-luciferase microRNA target expression vector (Promega, E1330, USA). Both mutant 3’UTR pmirGLO vectors were generated using the QuikChange II XL Site-Directed Mutagenesis Kit (Stratagene, USA) according to the manufacturer’s protocol. Next, 293 T cells were co-transfected with miR-214-3p mimic and either the VASH1 and MIEF2 pmirGLO vectors or the mutant VASH1 and MIEF2 3’UTR pmirGLO vectors. Then, luciferase activity was assessed using the Dual-Glo Luciferase Assay System (Promega, E2920).

Relative luciferase activity was calculated as the ratio of Firefly/Renilla luminescence.

### Quantitative polymerase chain reaction (qPCR)

Total RNA from left ventricular (LV) tissues and cells was extracted using Trizol (Invitrogen) and the EZ-press RNA Purification Kit (EZBioscience, B0004DP, China), respectively. The cDNA was synthesized with the Color Reverse Transcription Kit (EZBioscience, A0010CGQ). qPCR was performed using the SYBR Green qPCR Master Mix (EZBioscience, A0012-R2) and detected with the QuantStudio 7 Flex (Thermo Fisher Scientific). The transcript level of genes was calculated as 2^−△△Ct^, in which △Ct refers to the amplification cycles versus GAPDH, and the △△Ct refers to the △Ct versus the control group.

### Flow Cytometry Analysis

Cells were analyzed by flow cytometry as described previously ^[22]^. Briefly, the hEPs were collected after dissociation with Accutase for 5 min. The cells were fixed and permeabilized using the Staining Buffer Kit (Thermo Fisher Scientific) for 30 min, followed by incubation with the primary antibody for 1 h at room temperature (RT) and the PE-conjugated secondary antibody for 1 h at 4 °C. Cells were then analyzed on a flow cytometer (Beckman CytoFlex, USA). To assess apoptosis in hCMs, the Apoptosis Detection Kit (Yeasen, 40302ES60, China) was used. Results were quantified using FlowJo software.

### In vitro cytoprotection assay

The apoptosis of hCMs was induced by oxygen-glucose deprivation (OGD) conditions (glucose-free DMEM, 1% O_2_/5% CO_2_/94% N_2_). In brief, hCMs were cultured in four-chamber slides under normal or OGD conditions. Subsequently, PBS (control), exosomes, miR-214-3p agomir, or agomir NC were added, followed by a 48-h co-culture period. Apoptosis was assessed by staining cells with an In Situ Cell Death Detection Kit. Mitochondrial function and morphology were assessed using the JC-1 assay (Beyotime, C2006), Mito-Tracker Red CMXRos (Beyotime, C1035), and TEM.

### Immunofluorescent staining analysis

For in vitro immunofluorescent staining, cells were fixed with 4% paraformaldehyde (PFA), permeabilized with 0.1% Triton X-100 (Sigma, 9036-19-5), and then stained with Wilms tumor protein (WT1, Abcam, ab89901), zonula occludens 1 (ZO1, Millipore, 339100), β-Catenin (BD, 610154), vimentin (CST, 5741S), and α-smooth muscle actin (α-SMA, Abcam, ab5694). The frozen sections were brought to RT for 2 min and washed with PBS. Then the sections were fixed with 4% PFA, permeabilized with 0.4% Triton X-100, and stained with antibodies [CD31 (Abcam, ab28364), α-SMA (Sigma, a2549), and cardiac troponin T (cTnT, Abcam, ab8295)] and FITC-conjugated wheat germ agglutinin (WGA, Sigma, L4895). The Alexa 488- or 555-conjugated secondary antibodies were incubated for 2 h in the dark. DAPI was used for nuclear staining. To evaluate apoptosis post-MI, a one-step TUNEL Apoptosis Assay Kit (Beyotime, C1089) was used. Images were analyzed by ImageJ.

### Cell migration assay

The cell migration assay was performed as previously described ^[20]^. When HUVECs were grown to 100% confluence, then scratched and washed three times with PBS. Next, PBS (control) and exosomes were added to the culture media in the 12-well plates. The marked areas were photographed using a Zeiss inverted microscope before and after 8 h of different treatments.

### Tube formation assay

For the tube formation assay, 48-well plates were coated with 200 μL of Matrigel (Beyotime, C0382) per well and incubated at 37 °C for 30 min. HUVECs were seeded at 5×10^4^ cells/cm^2^ in basal medium, and PBS (control), exosomes, miR-214-3p agomir, or agomir NC were added to the culture medium. HUVECs were then maintained at 37 °C in humidified air with 5% CO_2_, and capillary-like structure formation was recorded using a Zeiss inverted microscope. Image J was used to quantify tube length ^[20]^.

### Measurement of adenosine triphosphate (ATP) levels

ATP production in cardiomyocytes was assessed using the ATP Assay Kit (Beyotime, S0026). Cardiomyocytes were seeded at 1 × 10^4^ cells per well in a 96-well plate and allowed to adhere for 24 h in a humidified incubator at 37°C with 5% CO_2_. After treatment, cells were lysed with the reagent, and the supernatant was collected. To measure ATP levels, 100 µL of the prepared ATP detection reagent was added to each well, and the plate was incubated at room temperature for 3-5 min to ensure complete ATP consumption. After incubation, luminescence was measured using a luminometer or similar light-measuring device. Luminescence values were recorded in relative light units (RLU), and ATP concentrations were calculated from a standard curve generated with known ATP concentrations.

### Measurement of reactive oxygen species (ROS) production

ROS production in cardiomyocytes was measured using the Reactive Oxygen Species Assay Kit (Beyotime, S0033S). Cardiomyocytes were seeded at 1 × 10^4^ cells per well in a 96-well plate and allowed to adhere for 24 h in a humidified incubator at 37°C with 5% CO_2_. After treatment, cells were incubated with 10 μM DCFH-DA in PBS for 20 min at 37°C. Cells were then washed three times with PBS to remove excess DCFH-DA. ROS production was measured using a fluorescence microplate reader with excitation and emission wavelengths of 488 nm and 525 nm, respectively.

### Isolation and culture of adult mouse cardiomyocytes (AMCMs)

Isolation of AMCMs was performed using an enzymatic method as previously described. ^[12]^ The resuspended cells were filtered and plated onto Matrigel-coated dishes. The plates were incubated in a humidified tissue culture incubator (37 °C, 5% CO_2_).

### Seahorse Metabolic Profiling

The Seahorse XF96 extracellular flux analyzer (Agilent Technologies) was used to assess mitochondrial function. The cells were seeded onto a 96-well Seahorse plate 48 h prior to the assay at a density of 1 × 10^4^ cells/well. One hour before the assay, the culture media were replaced with assay base media, and the plate was incubated at 37 °C in a non-CO_2_ incubator. The assay was then run according to the manufacturer’s instructions (Agilent). The collected data were analyzed with Agilent Wave software and graphed using GraphPad Prism 6.

### Analysis of Lactate Dehydrogenase (LDH) Activity

The sample preparation and LDH activity measurements were performed as previously reported ^[7]^. For cell samples, treat cells with Exo-N and Exo-H and set up controls. After treatment, aspirate the supernatant. Add 60 μL of LDH working solution to 120 μL of supernatant in a prepared 96-well plate. Mix well and incubate at RT in the dark for 30 min with gentle agitation on a horizontal shaker. The absorbance was recorded at 490 nm.

### Mouse MI model

All surgical procedures were performed in accordance with the Guidelines for Care and Use of Laboratory Animals, published by the US National Institutes of Health (NIH Publication, 8th Edition, 2011), and were approved by the Institutional Animal Care and Use Committee of the Shanghai Institutes for Biological Sciences. Mice aged 10–12 weeks were randomly divided into four groups: sham group, MI + PBS, MI + Exo-N, and MI + Exo-H. The mice were anesthetized and the MI model was induced by permanent ligation of the LAD coronary artery with an 8-0 Prolene suture, as previously described ^[23]^. The success of MI induction was further confirmed by a pale anterior wall and ST-segment elevation on the electrocardiogram. Exosomes mixed with PBS were then directly injected into the ischemic myocardium, and an equal volume of PBS was injected into the MI control group. The mice in the sham group underwent the same procedure, except for LAD ligation. Then the muscle and skin of the chest were sutured with a 6-0 Prolene suture. The body temperature was maintained at 37 °C during the surgical procedure. To determine the cardioprotective effects of miR-214-3p, animals subjected to MI were randomly assigned to the MI+miR-214-3p and MI+NC agomir groups, followed by intramyocardial injection of miR-214-3p agomir and NC agomir, respectively.

### Echocardiography

The primary outcome of this study will be cardiac function, assessed by echocardiography with a Vevo 2100 Imaging System (VisualSonics, Inc., Canada), as previously described ^[20]^. Briefly, the mice were lightly anesthetized with 2% isoflurane until the heart rate stabilized at 400–500 bpm; then, both conventional two-dimensional mode and M-mode images of the heart were acquired, and data on left ventricular ejection fraction (LVEF), left ventricular fractional shortening (LVFS), left ventricular end-systolic diameter (LVESD), and left ventricular end-diastolic diameter (LVEDD) were measured.

### Masson’s trichrome staining

For Masson’s trichrome-stained images, each heart was sectioned into five slices from the ligation point to the apex. The morphometric parameters, including total LV area and scar area, were analyzed blindly in the five heart slices. The scar size was calculated by dividing the total scar area by the LV area, as previously described.^[7]^

### Statistical analysis

Data are presented as mean ± standard error of mean (SEM). Firstly, the normality of all data was assessed using the Shapiro–Wilk normality test. Then, differences between the two mean values were evaluated via Student’s *t*-test, while analysis of variance (ANOVA) with Tukey’s post hoc test was used for multiple comparisons. All statistical analyses were performed using Statistical Product and Service Solutions (SPSS) software (version 18.0; IBM Corp., Armonk, NY, USA). *P* < 0.05 was considered statistically significant.

## Results

### Hypoxia-induced hEP activation and isolated exosome characterization

The differentiation of hEPs was performed as previously described (**Figure 1A**) ^[7, 18, 19]^. Immunofluorescent staining confirmed robust expression of the epicardial markers, WT1 and ZO1 (**Figure 1B**). Flow cytometry analysis further showed high differentiation efficiency, with 97.9 ± 0.53% of cells expressing WT1 (**Figure 1C**). Previous studies have shown that hypoxic conditioning induces EMT and enhances the paracrine function of primary epicardial cells ^[20, 24, 25]^. To determine whether hEPs exhibit comparable plasticity in response to hypoxia, hEPs were cultured under normoxic (hEP-N) or hypoxic (hEP-H) conditions (**Figure 1D**). Temporal gradient analysis of hypoxic exposure revealed a progressive decline in WT1 expression in hEPs, accompanied by time-dependent upregulation of EMT-related genes (**Figure S1A**). Notably, apoptosis in hEPs was significantly higher at 72 h than at 48 h (**Figure S1B**). Therefore, 48 h was selected as the optimal condition for subsequent experiments. Immunofluorescent staining of hEP-N and hEP-H also revealed that hypoxia upregulates snail family transcriptional repressor 2 (Slug) expression (a marker of EMT) and downregulates WT1 expression (**Figure 1E**), which is consistent with studies on primary EPs ^[9]^. Furthermore, qPCR analysis showed that hEP-H cells exhibited higher mRNA levels of EMT markers than hEP-N cells (**Figure 1F**). After 7 days of cultivation, additional markers consistent with fibroblasts and smooth muscle cells were evident in hEP-H cells, confirming their differentiation potential (**Figure 1G**).

**Figure 1.**
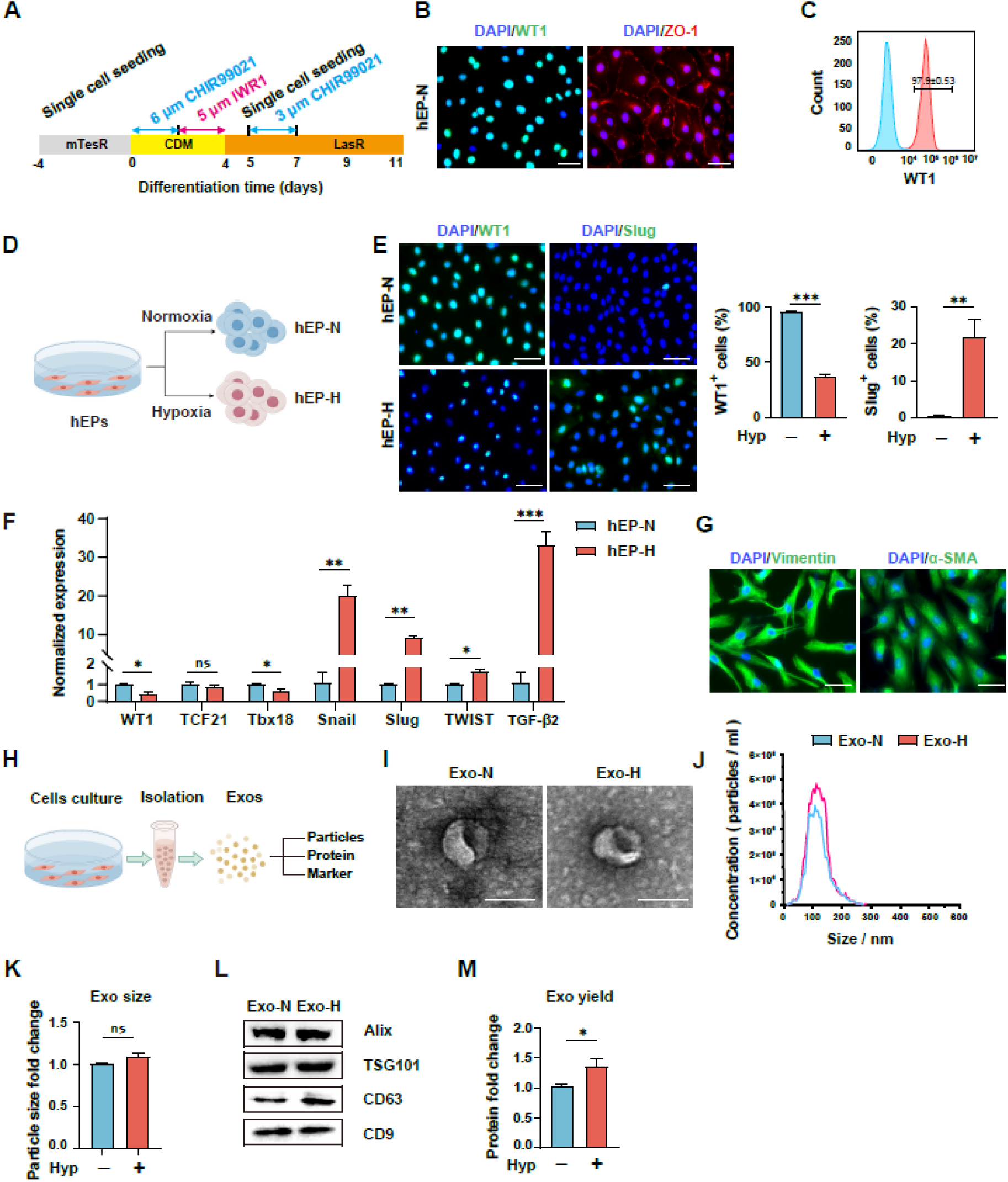
hiPSC-derived epicardial cells (hEPs) are activated in a hypoxic environment. **(A)** Schematic diagram of hEP differentiation. **(B)** Differentiated hEPs were analyzed by immunofluorescent staining for the hEP markers, WT1 and ZO-1. **(C)** Flow cytometry analysis of the differentiated cells for WT1 expression. **(D)** Schematic illustration of normoxia and hypoxia treatments. hEPs were divided into two experimental groups: normoxia-cultured hEPs (hEP-N, 21% O_2_) and hypoxia-cultured hEPs (hEP-H, 1% O_2_). **(E)** Representative immunofluorescent staining of hEP-N and hEP-H, along with quantification, revealed that the epithelial-mesenchymal transition (EMT) had occurred in hEPs (bar=50 μm). n=3. **(F)** Quantitative PCR (qPCR) analysis of representative EMT-related gene expression in hEP-N and hEP-H. n=3. **(G)** Representative immunofluorescent staining for vimentin and α-SMA in hEP-H after 7 days of cultivation (bar=50 μm). **(H)** Schematic illustration of the exosome isolation and characterization process. Exosomes isolated from hEP-N and hEP-H were designated as Exo-N and Exo-H, respectively. **(I)** Representative transmission electron microscope (TEM) images of Exo-N and Exo-H (bar=100 nm)**. (J)** Size distributions of Exo-N and Exo-H, as measured by nanoparticle tracking analysis (NTA). **(K)** Particle sizes of Exo-N and Exo-H. **(L)** Western blot analysis of exosome markers (Alix, TSG101, CD63, and CD9) in exosomes. **(M)** Exosome yields across different groups, as determined by the exosome protein/cell protein ratio. Quantitative data are expressed as mean ± SEM. Comparisons between two groups were analyzed using a two-tailed Student’s *t*-test (E, F, K, and M). \*\*\**P* < 0.001, \*\**P* < 0.01, \**P* < 0.05, and ^ns^*P* ≥ 0.05. WT1: Wilms tumor protein, ZO-1: zonula occludens-1, α-SMA: α-smooth muscle actin, Alix: ALG-2 interacting protein-X, TSG101: tumor susceptibility gene 101, CD63: cluster of differentiation 63, and CD9: cluster of differentiation 9.

Given the observed phenotypic divergence between hEP-N and hEP-H, we then examined the differences in their paracrine capabilities. Exosomes were isolated from the conditioned media of hEP-N (Exo-N) and hEP-H (Exo-H) by ultracentrifugation as described previously (**Figure 1H**) ^[12, 20]^. Both types of exosomes exhibited the typical double-membrane-bound, cup-shaped morphology (**Figure 1I**). Nanoparticle tracking analysis (NTA) revealed that the mean size of the secreted exosomes was approximately 100-110 nm (**Figure 1J and K**). Western blot analysis further confirmed the presence of exosome marker proteins (ALIX, TSG101, CD63, and CD9) in the Exo-N and Exo-H samples (**Figure 1L**). However, the exosome yield in the hypoxic group was higher than in the normoxic group (**Figure 1M**). These data collectively demonstrate that exosomes from both hEP-N and hEP-H were successfully isolated and that hypoxia stimulates hEPs to secrete more exosomes.

### hEP-Exos enhance the angiogenic potential of ECs and attenuate cardiomyocyte apoptosis in vitro

Exo-N and Exo-H were resuspended in PBS and added to HUVECs for functional assays (**Figure 2A**). Fluorescent labeling of hEP-Exos with the lipophilic dye DiI showed efficient cellular internalization by HUVECs (**Figure 2B**). HUVECs migration assays revealed enhanced migratory capacity in cells treated with Exo-N or Exo-H, with Exo-H producing more pronounced effects (**Figure 2C**). Similarly, treatment with Exo-N or Exo-H markedly increased the number of junctions and total branching length in HUVECs compared with the control group, with Exo-H exhibiting greater efficacy than Exo-N (**Figure 2D**). Subsequently, we investigated the effects of Exo-N and Exo-H on cardiomyocytes (**Figure 2E**). Fluorescent labeling of hEP-Exos with the lipophilic dye DiI showed efficient cellular internalization by hCMs (**Figure 2F**). Immunofluorescent staining showed that both Exo-N and Exo-H attenuated hCM apoptosis (TUNEL^+^) under OGD injury, with Exo-H exhibiting a more pronounced therapeutic effect (**Figure 2G**).

**Figure 2.**
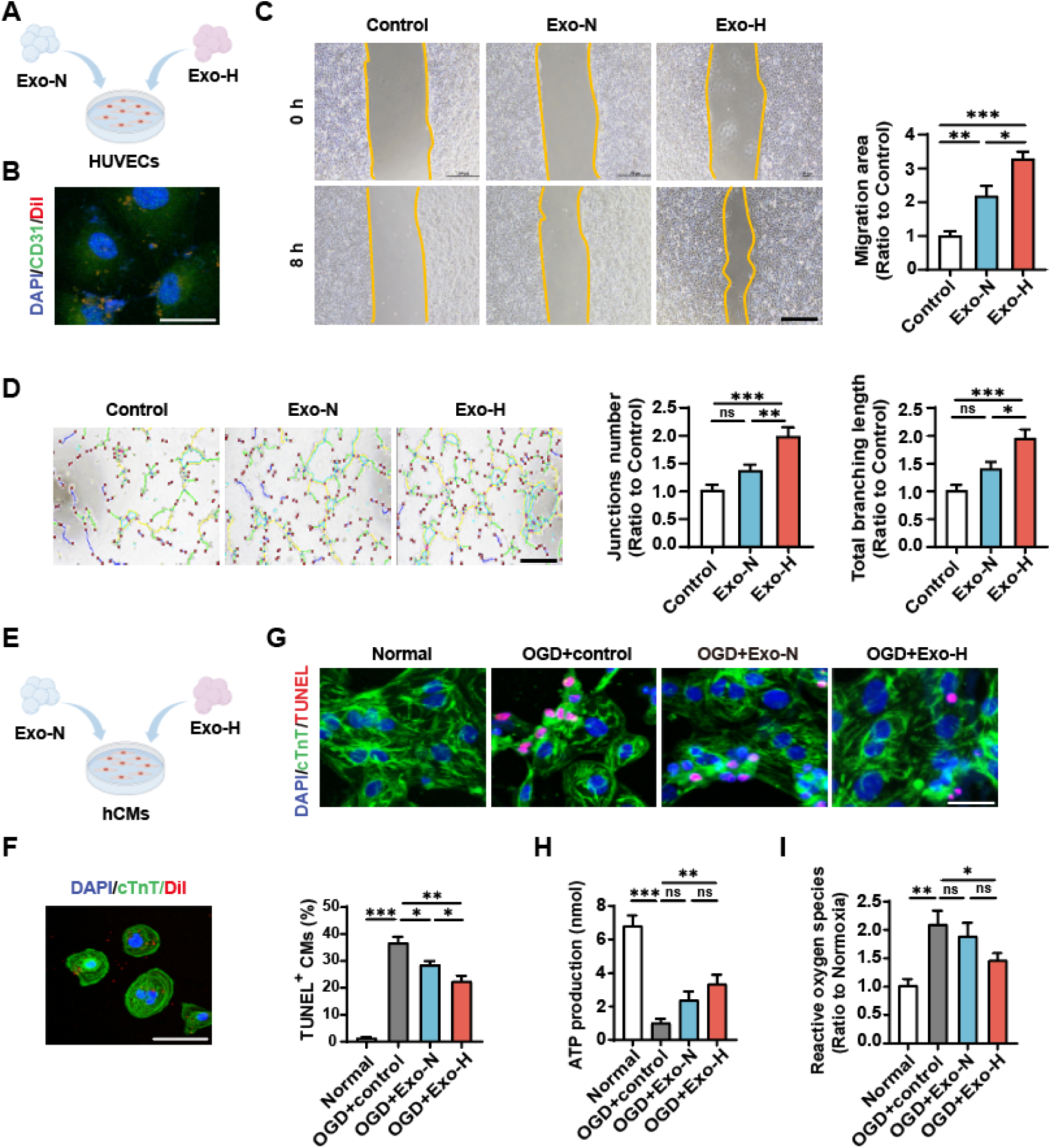
In vitro effects of Exo-H and Exo-N on endothelial cells and cardiomyocytes. **(A)** Schematic illustration of the treatment of human umbilical vein endothelial cells (HUVECs) with Exo-N and Exo-H. **(B)** Immunofluorescent staining revealed that HUVECs successfully internalized DiI-labeled exosomes. **(C)** Migration assessment of HUVECs treated with Exo-N and Exo-H (bar=100 μm). n=5. **(D)** Tube formation assessment of HUVECs treated with Exo-N and Exo-H (bar=200 μm). n=5. **(E)** Schematic illustration of hiPSC-derived cardiomyocytes (hCMs) treated with Exo-N and Exo-H. **(F)** Immunofluorescent staining revealed that hCMs successfully internalized DiI-labeled exosomes. **(G)** Representative images of TUNEL^+^ hCMs and quantitative percentages of TUNEL^+^ cells after oxygen-glucose deprivation (OGD) injury (bar=50 μm). **(H)** Adenosine triphosphate (ATP) production in hCMs following OGD injury. **(I)** Reactive oxygen species (ROS) release in hCMs after OGD injury. Quantitative data are presented as mean ± SEM. For comparisons involving more than two groups, a one-way ANOVA followed by Tukey’s post-hoc test was performed to determine statistical significance between specific experimental conditions (C, D, F, H, and I). \*\*\**P* < 0.001, \*\**P* < 0.01, \**P* < 0.05, and ^ns^*P* ≥ 0.05.

Mitochondria, the primary organelles responsible for energy metabolism in cardiomyocytes, undergo significant changes in metabolic activity under stress. These changes can, in turn, influence cardiomyocyte apoptosis by disrupting intracellular energy homeostasis ^[26]^. Building on this foundation, we further investigate the dynamic characteristics of energy metabolism in hCMs under OGD conditions. Quantitative analyses showed that OGD injury markedly decreased adenosine triphosphate (ATP) production while increasing ROS generation. Notably, both Exo-N and Exo-H treatments can counteract these pathological alterations (**Figure 2H and I**). Consistent with this, JC-1 staining confirmed that they can also improve mitochondrial membrane potential in cardiomyocytes after OGD injury (**Figure S2A**). Notably, Exo-H treatment showed better effects. These findings demonstrate the mitochondrial-protective capacity of hEP-Exos to mitigate cardiomyocyte apoptosis under ischemic conditions, with Exo-H exhibiting superior efficacy.

### Intramyocardial delivery of hEP-Exos improves cardiac function and reduces scar size in mice with MI

To evaluate the therapeutic efficacy of exosomes delivered during the acute phase of MI, Exo-N and Exo-H were injected intramyocardially into mice with acute MI (**Figure 3A**). Both the Exo-N and Exo-H groups exhibited significantly higher survival rates than the control group (**Figure 3B**). Cardiac function was assessed via echocardiography at 2-, 7-, 14-, and 28-days after MI or sham surgery. On day 2 post-MI, changes in LVEF, LVFS, LVESD, and LVEDD were comparable across the PBS, Exo-N, and Exo-H groups, suggesting a similar extent of initial infarction. However, over the subsequent 28-day observation period, LVEF, LVFS, LVESD, and LVEDD, which were significantly worsened in the PBS group, showed marked improvement in both the Exo-N and Exo-H groups. Importantly, the Exo-H group showed a superior recovery trend compared to the Exo-N group (**Figure 3C**). This functional improvement was further corroborated by Masson’s trichrome staining, which showed that the scar area relative to LV area was significantly smaller in the Exo-N and Exo-H groups than in the PBS group at day 28 post-MI (**Figure 3D**). Additionally, adverse LV remodeling, evidenced by an increased heart-to-body weight ratio and larger cardiomyocyte cross-sectional area in the border zone, was significantly mitigated by Exo-N and Exo-H treatments, with Exo-H exhibiting the more pronounced effects (**Figure 3E and F**).

**Figure 3.**
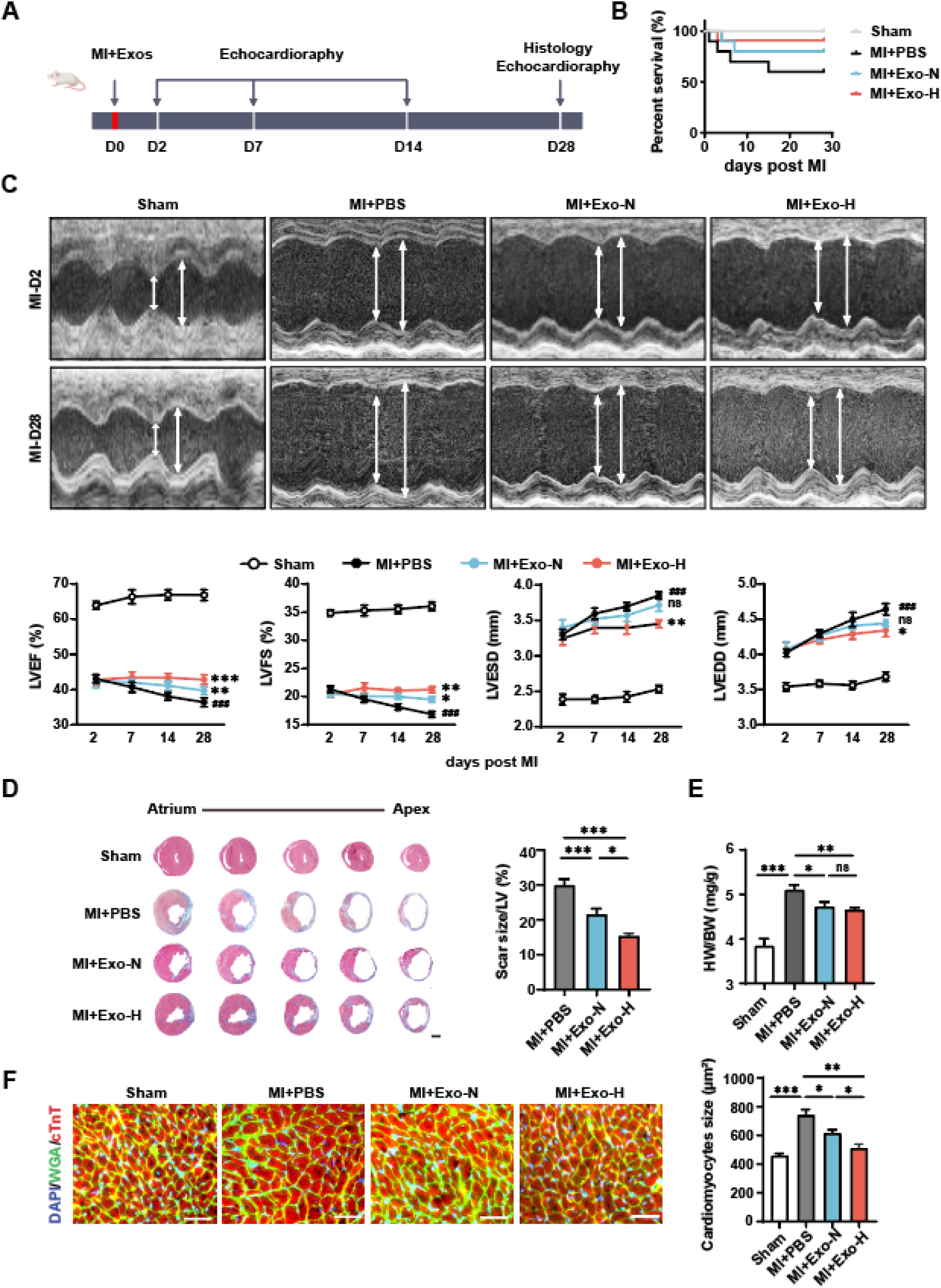
Cardioprotective effects of Exo-H and Exo-N delivered to acutely infarcted mouse hearts. **(A)** Myocardial infarction (MI) was surgically induced in mice via ligation of the left anterior descending (LAD) coronary artery on day 0 (D0), followed immediately by administration of exosomes. Serial echocardiography was performed on days 2, 7, 14, and 28 after MI. Terminal tissue collection was performed on day 28 (D28) for comprehensive histological evaluation. **(B)** Kaplan–Meier curve showing mortality after MI. n=15 mice per group. **(C)** Echocardiographic analysis of left ventricular ejection fraction (LVEF), left ventricular fractional shortening (LVFS), left ventricular end-systolic diameter (LVESD), and left ventricular end-diastolic diameter (LVEDD). n=8 mice per group. \*\*\**P* < 0.001, \*\**P* < 0.01, \**P* < 0.05, and ^ns^*P* ≥ 0.05 versus the MI group; ^###^*P* < 0.001 versus the sham group. **(D)** Representative cross-sectional images and quantitative analysis of scar size stained with Masson’s trichrome on day 28 after MI (bar=1 mm). n=5. **(E)** Heart-to-body weight ratio 28 days after MI. n=5. **(F)** Sections from the border zone (BZ) of the infarcted heart were stained with wheat germ agglutinin (WGA) to identify the cellular borders and with cardiac troponin T (cTnT) antibodies to visualize cardiomyocytes; nuclei were counterstained with DAPI (bar=100 μm). n=5. Quantitative data are presented as mean ± SEM. For echocardiographic data, a two-way repeated-measures ANOVA followed by Bonferroni’s post hoc test was used to compare differences between groups over time (C). Scar size and histological quantifications (D, E, and F) were analyzed using one-way ANOVA followed by Tukey’s multiple comparison test. \*\*\**P* < 0.001, \*\**P* < 0.01, \**P* < 0.05, and ^ns^*P* ≥ 0.05.

### hEP-Exos enhance angiogenesis and attenuate cardiomyocyte apoptosis in infarcted mouse hearts

Angiogenesis plays a critical role in reducing cell death and limiting fibrosis in infarcted hearts. To investigate the pro-angiogenic effects of hEP-Exos, we examined blood vessel density in the infarcted hearts. Immunofluorescent staining revealed that the number of α-SMA⁺ vessels (**Figure 4A**) and CD31⁺ vessels (**Figure 4B**) at the border zone and infarct zone of infarcted hearts on day 28 post-MI was significantly higher in the Exo-N and Exo-H groups than in the PBS group. Furthermore, the number of TUNEL⁺ cardiomyocytes in the border zone of infarcted hearts after MI was significantly reduced in the Exo-N and Exo-H groups compared to the PBS group (**Figure 4C**). Notably, the Exo-H group exhibited a greater effect than the Exo-N group (**Figure 4A-C**), indicating that exosomes derived from hEP-H provide superior protection in enhancing angiogenesis and reducing cardiomyocyte death. In vivo observations of metabolic dysfunction prompted further investigation using AMCMs isolated from MI hearts. Seahorse metabolic analysis demonstrated that MI significantly reduced the oxygen consumption rate (OCR) in AMCMs, whereas exosome treatment restored OCR. This metabolic improvement, with increased ATP production and decreased ROS release, correlated with reduced cardiomyocyte apoptosis, and Exo-H showed superior therapeutic efficacy compared to Exo-N (**Figure 4D-F**).

**Figure 4.**
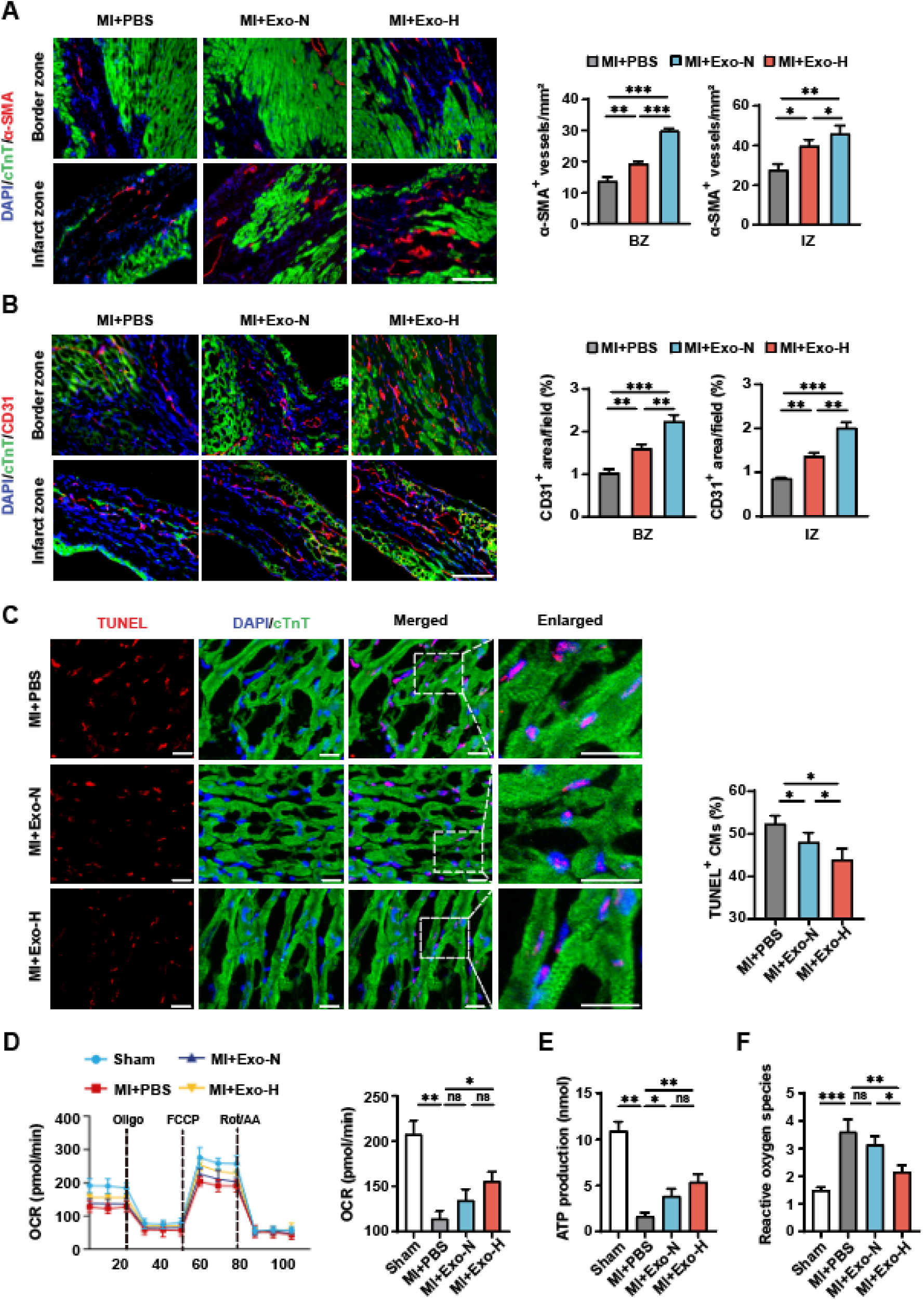
In vivo effects of Exo-H versus Exo-N on endothelial cells and cardiomyocytes. **(A)** Representative images and quantitative data from immunofluorescent staining for α-SMA^+^ smooth muscle cells in the BZ and the infarct zone (IZ) (bar=100 μm). n=5. **(B)** Representative images and quantitative data from immunofluorescent staining for CD31^+^ endothelial cells in the BZ and IZ (bar=100 μm). n=5. **(C)** Representative images and quantification of immunofluorescent staining for TUNEL^+^ cardiomyocytes in infarcted hearts (bar=20 μm). n=5. **(D)** Seahorse assays in adult mouse cardiomyocytes (AMCMs) post-MI. n=3. **(E)** The ATP production in AMCMs after MI. n=3. **(F)** The release of ROS in AMCMs after MI. n=3. Quantitative data are presented as mean ± SEM. For experiments involving three or more groups (A, B, C, D, E, and F), a one-way ANOVA followed by Tukey’s post-hoc test for multiple comparisons was applied. \*\*\**P* < 0.001, \*\**P* < 0.01, \**P* < 0.05, ^ns^*P* ≥ 0.05. α-SMA: alpha smooth muscle actin, CD31: platelet endothelial cell adhesion molecule-1.

### Exo-H exhibits significant enrichment of miR-214-3p compared to Exo-N

Exosomes from a wide range of sources have been shown to carry a myriad of cargoes, including miRNAs that have been implicated as potential mediators of regenerative signals in the context of cardiovascular disease ^[12, 13]^. In our findings above, Exo-H demonstrated superior cardioprotective effects compared to Exo-N. To investigate the underlying molecular basis, we extracted total RNA from Exo-N and Exo-H and performed miRNA sequencing. The results identified 255 miRNAs exclusively present in Exo-N and 87 miRNAs exclusively present in Exo-H. Additionally, 115 miRNAs were significantly upregulated in Exo-H compared with Exo-N, with miR-214-3p showing the most pronounced increase.

### miR-214-3p promotes angiogenesis through suppressing VASH1

Integrated bioinformatics analysis utilizing PITA, miRmap, and microT databases identified 651 putative target genes of miR-214-3p through cross-algorithm consensus (**Figure 5A**). Subsequent intersection analysis between the predicted target genes and angiogenesis-related genes curated from GeneCards identified six candidate genes (**Figure 5B**). qPCR analysis of mRNA in HUVECs revealed VASH1 as the most significantly downregulated gene among the candidate genes (**Figure 5C**). VASH1 was reported as a negative feedback regulator of angiogenesis. To verify the interaction between VASH1 and miR-214-3p, a dual-luciferase reporter assay was performed using plasmids containing either the wild-type (WT) or mutant VASH1 3’-UTR sequence (**Figure 5D**). Luciferase activity analysis revealed a significant reduction in activity in the VASH1 3’-UTR WT group under miR-214-3p treatment, whereas this effect was abolished in the VASH1 3’-UTR mutant group (**Figure 5E**). To elucidate the critical role of miR-214-3p in angiogenesis, gain- and loss-of-function experiments in vitro were conducted. Tube formation assays showed that the enhanced tube-forming capacity of ECs in the Exo-H^NC^ group was abolished in the Exo-H^anti-miR-214-3p^ group and reproduced in the miR-214-3p agomir group (**Figure 5F**). Western blot analysis revealed a marked reduction in VASH1 protein levels and a corresponding increase in VEGFA protein levels in the Exo-H^NC^ group. These changes were reversed in the Exo-H^anti−miR−214−3p^ group and recapitulated in the miR-214-3p agomir group (**Figure 5G**).

**Figure 5.**
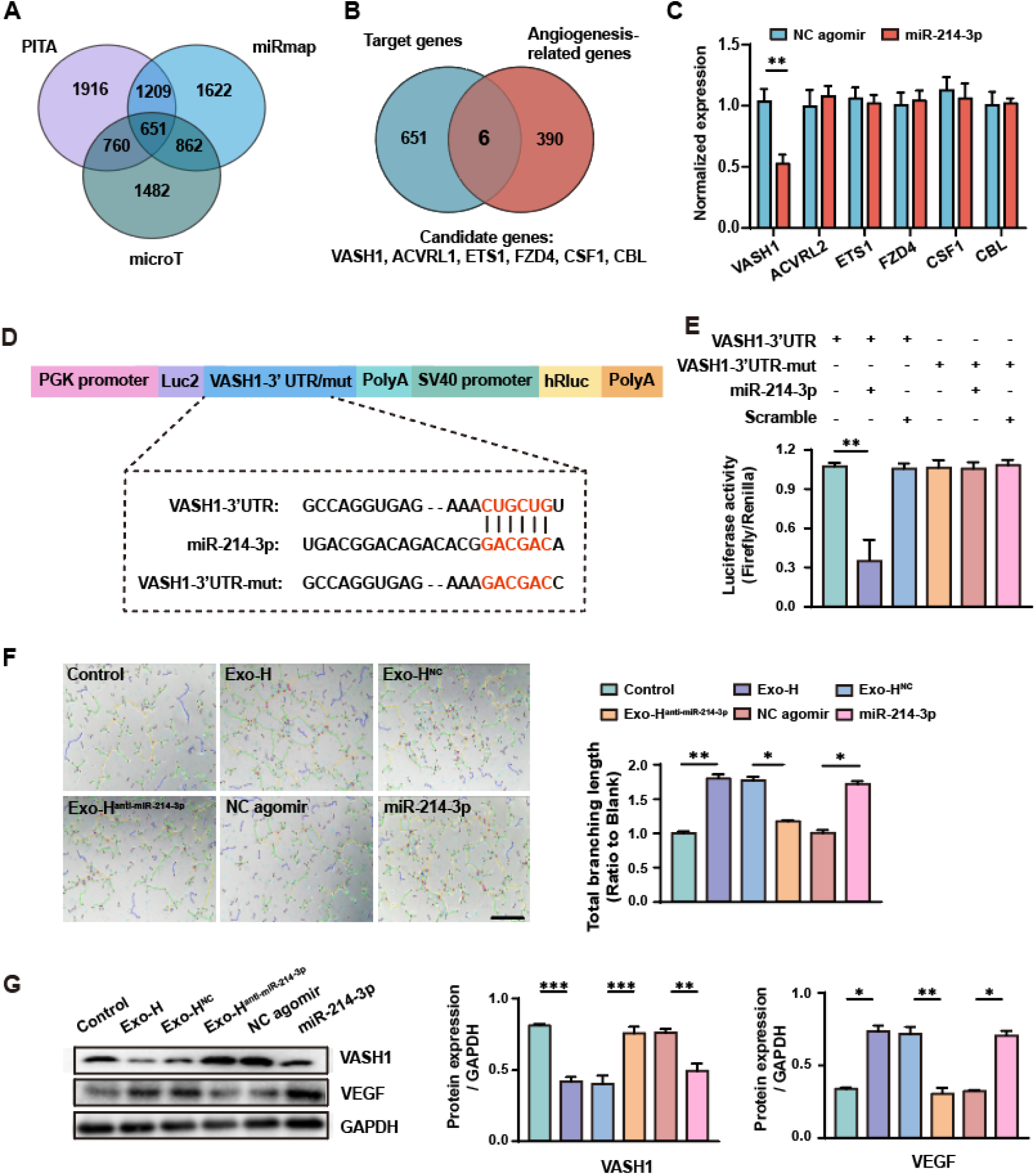
miR-214-3p promotes angiogenesis by suppressing VASH1. **(A)** Bioinformatic analysis using the PITA, miRMAP, and microT algorithms predicted that 651 genes are downstream of miR-214-3p. **(B)** Vascular-related genes were retrieved from GeneCards. Subsequent intersection with our predicted targets identified 6 candidate genes: VASH1, ACVRL1, ETS1, FZD4, CSF1, and CBL. **(C)** qPCR was performed to assess mRNA expression levels of the six candidate genes in endothelial cells. **(D)** Wild-type or mutant dual-luciferase reporter plasmids were constructed according to the predicted binding sequence in the 3’ UTR of VASH1 or the mutant sequence. **(E)** The activities of Renilla and Firefly luciferase were determined using the Dual-Luciferase Reporter Assay System. **(F)** Tube formation assessment of HUVECs treated with Exo-H, Exo-H^NC^, Exo-H^anti-miR-214-3p^, agomir NC, and miR-214-3p agomir (bar=100 mm). n=3. **(G)** Representative images and quantification of Western blots of VASH1 and VEGF. Quantitative data are presented as mean ± SEM. For comparisons between two independent groups (C), a two-tailed Student’s *t*-test was used. For experiments involving three or more groups (E, F, and G), a one-way ANOVA followed by Tukey’s post-hoc test for multiple comparisons was applied. \*\*\**P* < 0.001, \*\**P* < 0.01, \**P* < 0.05, and ^ns^*P* ≥ 0.05. VASH1: Vasohibin 1, ACVRL1: activin A receptor-like type 1, ETS1: ETS proto-oncogene 1, FZD4: frizzled class receptor 4, CSF1: colony-stimulating factor 1, and CBL: Cbl proto-oncogene.

### miR-214-3p reduces cardiomyocyte apoptosis through suppressing MIEF2

Building on our above findings of improved mitochondrial function, we performed an integrative genomic analysis to elucidate the mechanistic link among hEP-Exo-mediated regulation, mitochondrial function, and cardiomyocyte apoptosis. By systematically intersecting predicted target genes with mitochondrial-associated and apoptosis-related genes (GeneCards), we identified four high-probability candidate genes that may mediate protective effects (**Figure 6A**). qPCR analysis of mRNA in hCMs revealed MIEF2 as the most significantly downregulated gene among the candidate genes (**Figure 6B**). MIEF2 encodes an outer mitochondrial membrane protein that plays a crucial role in regulating mitochondrial morphology by directly recruiting the fission mediator Drp1 to the mitochondrial surface ^[27]^. To verify the interaction between MIEF2 and miR-214-3p, a dual-luciferase reporter assay was performed using plasmids containing either the WT or mutant MIEF2 3’-UTR sequence (**Figure 6C**). Luciferase activity analysis revealed a significant reduction in activity in the MIEF2 3’-UTR WT group under miR-214-3p treatment, whereas this effect was abolished in the MIEF2 3’-UTR mutant group (**Figure 6D**). To elucidate the critical role of miR-214-3p in cardiomyocyte apoptosis, we conducted gain- and loss-of-function experiments in vitro. TUNEL assays and LDH release showed that the decreased cardiomyocyte apoptosis in the Exo-H^NC^ group was abolished in the Exo-H^anti-miR-214-3p^ group and reproduced in the miR-214-3p agomir group (**Figure 6E and F**). Western blot analysis revealed a marked reduction in MIEF2 protein levels in the Exo-H^NC^ group. These changes were reversed in the Exo-H^anti−miR−214−3p^ group and recapitulated in the miR-214-3p agomir group (**Figure 6G**).

**Figure 6.**
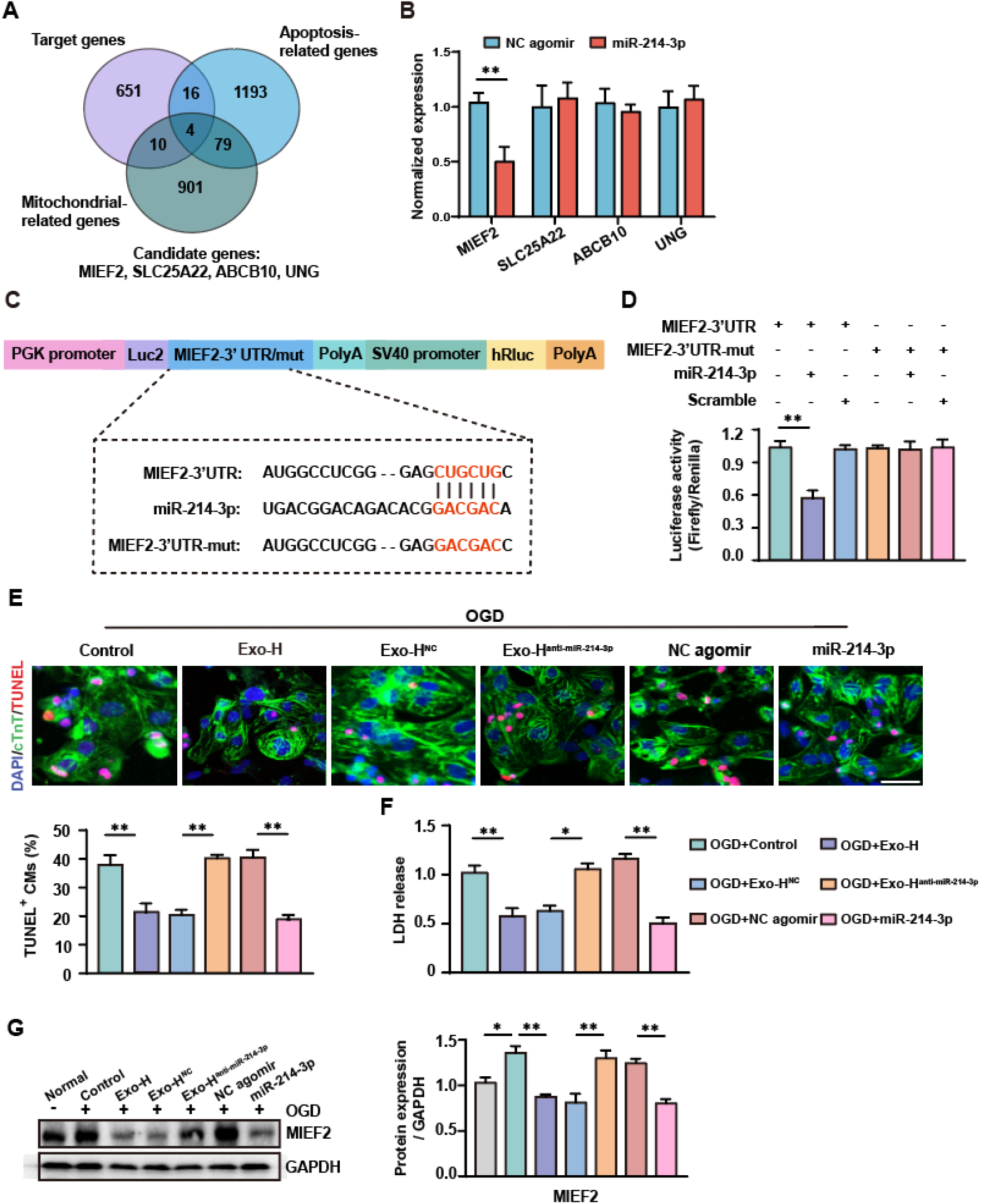
miR-214-3p reduces cardiomyocyte apoptosis by suppressing MIEF2. **(A)** Apoptosis- and mitochondrial-related genes were retrieved from the GeneCards database. Subsequent intersection with our predicted targets identified 4 candidate genes: MIEF2, SLC25A22, ABCB10, and UNG. **(B)** qPCR was performed to assess mRNA expression of the four candidate genes in endothelial cells. **(C)** Wild-type or mutant dual-luciferase reporter plasmids were constructed according to the predicted binding sequence in the 3’ UTR of MIEF2 or the mutant sequence. **(D)** The activities of Renilla and Firefly luciferase were determined using the Dual-Luciferase Reporter Assay System. **(E)** Representative images and quantification of immunofluorescent staining for TUNEL^+^ cardiomyocytes (bar=20 μm). n=3. **(F)** LDH release in the supernatant after OGD. **(G)** Representative images and quantification of Western blots of MIEF2. Quantitative data are presented as mean ± SEM. For comparisons between two independent groups (B), a two-tailed Student’s *t*-test was used. For experiments involving three or more groups (D, E, F, and G), a one-way ANOVA followed by Tukey’s post-hoc test for multiple comparisons was applied. \*\**P* < 0.01 and \**P* < 0.05. MIEF2: mitochondrial elongation factor 2, SLC25A22: solute carrier family 25 member 22, ABCB10: ATP binding cassette subfamily B member 10, and UNG: uracil DNA glycosylase.

### miR-214-3p attenuates cardiomyocyte mitochondrial fission

As the organ with the highest energy demand in the body, the heart relies heavily on mitochondria to meet its substantial energy requirements and support cardiac excitation–contraction coupling. TEM examination of mitochondrial morphology in hCMs showed that OGD injury significantly increased mitochondrial fission in cardiomyocytes, but that this pathological process was effectively reversed by miR-214-3p agomir. (**Figure 7A**). Western blot analysis revealed distinct patterns of Drp1 subcellular localization across different treatment conditions. Although total cellular Drp1 levels remained unchanged after OGD injury, a significant increase in mitochondrial Drp1 accumulation was observed. Treatments with Exo-H, Exo-H^NC^, and miR-214-3p agomir effectively reduced mitochondrial Drp1 levels, whereas control, agomir NC, and Exo-H^anti-miR-214-3p^ treatments showed no significant effects (**Figure 7B**). To further investigate mitochondrial fission dynamics, cardiomyocytes were examined using a mitochondrial probe. A marked increase in mitochondrial fission was observed in cardiomyocytes after OGD injury, but this increase was significantly alleviated in groups treated with Exo-H, Exo-H^NC^, and the miR-214-3p agomir. Interestingly, the protective effects were abolished upon miR-214-3p knockdown in the Exo-H group (**Figure 7C**). Furthermore, JC-1 staining corroborated these findings, showing that loss of miR-214-3p abrogated Exo-H’s ability to mitigate the decline in mitochondrial membrane potential, whereas miR-214-3p agomir reproduced these effects (**Figure 7D**). Additionally, the assessment of ATP and ROS production followed a similar pattern, further supporting the protective role of Exo-H and miR-214-3p agomir in mitigating mitochondrial dysfunction (**Figure 7E and F**).

**Figure 7.**
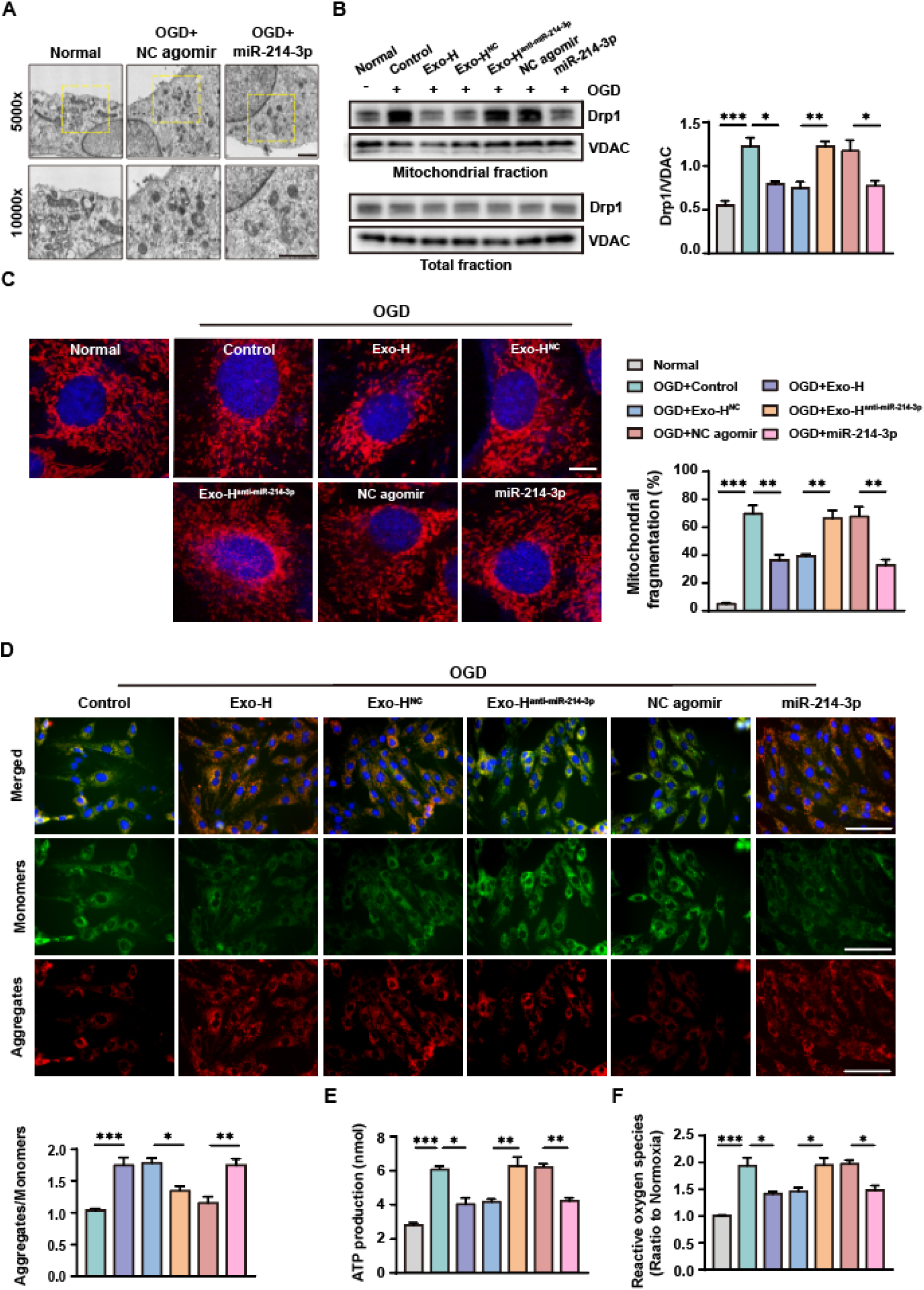
miR-214-3p reduces mitochondrial fission in cardiomyocytes with OGD injury. **(A)** Mitochondrial morphology in hCMs was analyzed using TEM. Mitochondrial size and arrangement are shown at 5,000x and 10,000x magnification (bar=2 μm). **(B)** Western blot analysis of Drp1 and MIEF2 expression in total and mitochondrial fractions. Voltage-dependent anion channel (VDAC) served as a loading control. Quantification of Drp1 and MIEF2 levels is shown in the histogram. **(C)** Mitochondria in hCMs from different groups after OGD injury were labeled with Mito-Tracker Red CMXRos to quantify the number of cells with mitochondrial fragmentation (bar = 10 μm). **(D)** Following OGD, hCMs were stained with JC-1. The degree of mitochondrial depolarization was assessed by calculating the red-to-green fluorescence ratio (bar = 100 μm). n=3. **(E)** ATP production in hCMs after OGD injury. n=3. **(F)** ROS release in hCMs after OGD injury. n=3. Quantitative data are presented as mean ± SEM. For experiments involving three or more groups (B, C, D, E, and F), a one-way ANOVA followed by Tukey’s post-hoc test for multiple comparisons was applied. \*\*\**P* < 0.001, \*\**P* < 0.01, and \**P* < 0.05.

### Intramyocardial delivery of miR-214-3p improves post-MI cardiac function and reduces scar size

To evaluate the therapeutic efficacy of miR-214-3p delivered during the acute phase of MI, agomir NC, and miR-214-3p agomir were intramyocardially injected into mice with acute MI (**Figure 8A**). The miR-214-3p agomir group exhibited significantly improved survival compared to the agomir NC and control groups (**Figure 8B**). Furthermore, echocardiographic analysis and Masson’s trichrome staining revealed substantial improvements in cardiac function and a reduction in fibrosis in the miR-214-3p agomir group (**Figure 8C and D**). We next examined the effects of miR-214-3p on angiogenesis and anti-apoptosis in vivo. Immunofluorescent staining demonstrated that miR-214-3p treatment significantly increased CD31^+^ vascular density in both the border zone and infarct zone, while reducing TUNEL^+^ cardiomyocytes in post-MI mouse hearts (**Figure 8E and F**). TEM analysis revealed a significant increase in mitochondrial fission in the agomir NC group, whereas miR-214-3p treatment effectively reversed this pathological process (**Figure 8G**). Complementary Seahorse metabolic assays further demonstrated that miR-214-3p administration substantially improved mitochondrial functional parameters (**Figure 8H**), with increased ATP production and decreased ROS release being notably observed (**Figure S3A and B**). These results demonstrate that miR-214-3p exhibits therapeutic effects comparable to Exo-H, and that Exo-H mediates its cardioprotective effects at least partially through miR-214-3p.

**Figure 8.**
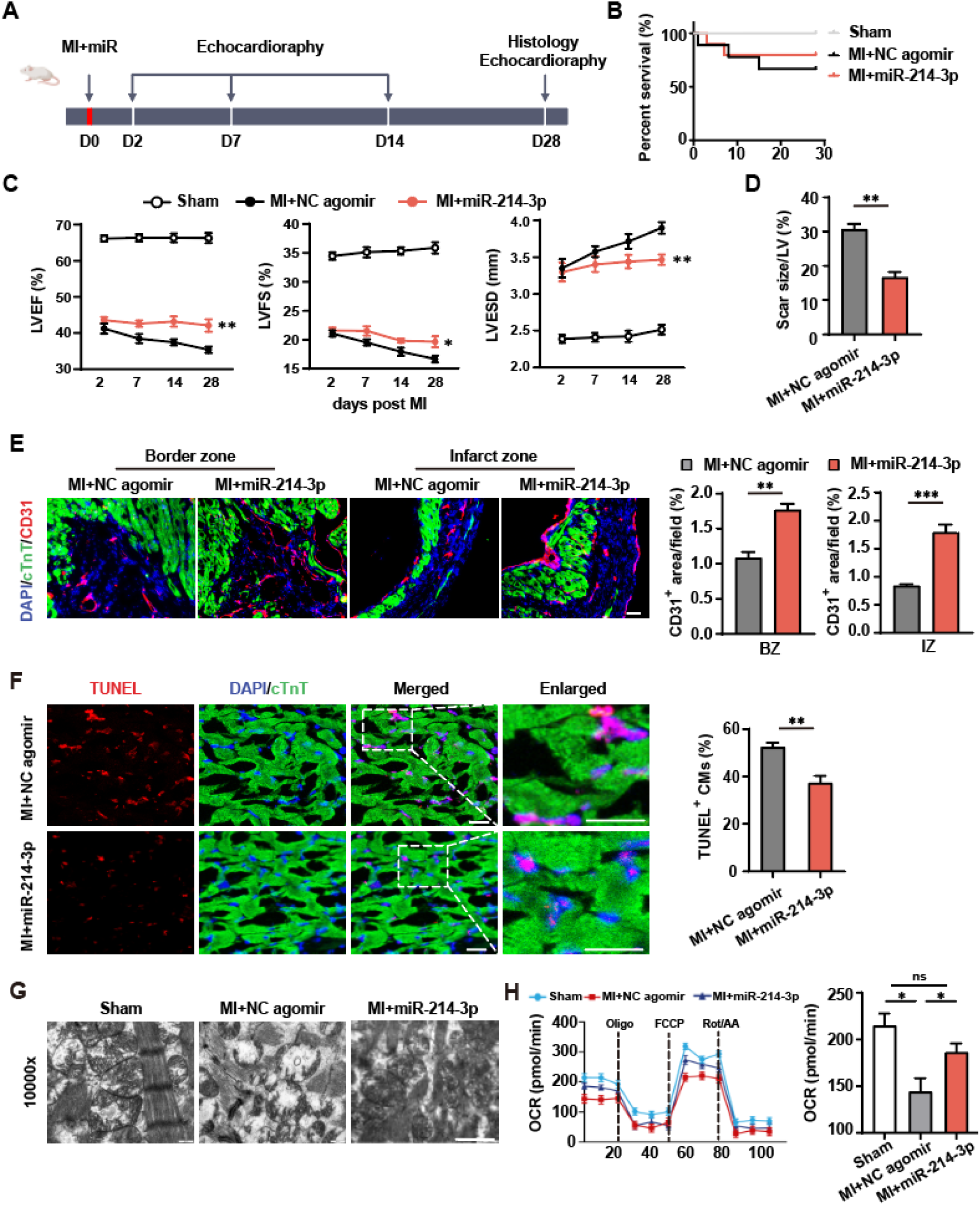
Administration of miR-214-3p effectively alleviates cardiac damage in mice with MI. **(A)** Schematic showing the treatment and analysis process involving the use of miR-214-3p agomir and agomir NC. **(B)** Kaplan-Meier curve showing mortality across groups after MI or sham operation. n=15 mice per group. **(C)** Echocardiographic analysis of LVEF, LVFS, and LVESD. n=8 mice per group. \*\**P* < 0.01 and \**P* < 0.05 versus the agomir NC group; ^###^*P* < 0.001 versus the sham group. **(D)** Quantitative analysis of scar size stained with Masson’s trichrome on day 28 post-MI (bar=1 mm). n=5. **(E)** Representative images and quantitative data of immunofluorescent staining for CD31^+^ endothelial cells in the BZ and IZ (bar=100 μm). n=5. **(F)** Representative images and quantification of immunofluorescent staining for TUNEL^+^ cardiomyocytes in infarcted hearts (bar=20 μm). n = 3. **(G)** Mitochondrial fission in the heart was examined by TEM. **(H)** Seahorse assays on AMCMs post-MI. n=3. Quantitative data are presented as mean ± SEM. For echocardiographic data (C), a two-way repeated measures ANOVA followed by Bonferroni’s post-hoc test was used to compare differences between groups over time. For comparisons between two independent groups (D, E, and F), a two-tailed Student’s *t*-test was applied. For comparisons of three or more groups (H), one-way ANOVA followed by Tukey’s multiple comparison test was used. \*\*\**P* < 0.001, \*\**P* < 0.01, \**P* < 0.05, and ^ns^*P* ≥ 0.05.

## Discussion

Cardiovascular regeneration remains a central challenge in the treatment of ischemic heart disease. Despite advances in pharmacological and interventional therapies, the capacity of the adult mammalian heart for self-repair is limited, and MI continues to be a leading cause of morbidity and mortality worldwide. In this study, we propose a novel acellular therapeutic strategy utilizing hEP-Exos, which is further enhanced by hypoxic preconditioning, to confer greater regenerative potency. Our findings reveal that Exo-H exerts its therapeutic effects through multiple converging mechanisms, including stimulating angiogenesis and inhibiting cardiomyocyte apoptosis by preserving mitochondrial integrity. These actions are orchestrated in part by miR-214-3p, a microRNA enriched within Exo-H, which simultaneously targets VASH1 (a known anti-angiogenic factor) and MIEF2 (a critical regulator of mitochondrial fission). This dual-targeting mechanism represents a finely tuned approach to repairing the infarcted myocardium, addressing the two essential but often independently targeted processes of vascular regeneration and cardiomyocyte survival in cardiac repair. These findings build on existing knowledge and shed light on the mechanisms underlying the cardioprotective effects of hEP-Exos, suggesting that these conditioned cells may be an effective acellular therapy for treating MI.

Epicardial cells, which are derived from the proepicardium, play a crucial role in both cardiac development and the response to injury. During development, these cells undergo EMT, giving rise to epicardial-derived cells (EPDCs), which can differentiate into fibroblasts, coronary smooth muscle cells, and pericytes.^[3]^ They promote vascularization by secreting angiogenic factors, such as VEGF, and influence myocardial growth through secreting fibroblast growth factors (FGFs) and retinoic acid. ^[28]^ Following cardiac injury, epicardial cells reactivate and undergo EMT again, generating EPDCs that participate in the formation of scar tissue. These cells secrete growth factors, including insulin-like growth factor 1 (IGF-1) and hepatocyte growth factor (HGF), which modulate the inflammatory response, promote angiogenesis, and support cardiomyocyte regeneration ^[29]^. Despite their stabilizing role in the injured heart, their limited proportion in the heart results in restricted therapeutic efficacy. This limitation is a key focus of therapeutic research aimed at improving cardiac repair. hESC-derived EPs replicate this reparative potential in vitro ^[7]^. Notably, here, we demonstrate that culturing hEPs under hypoxic conditions not only triggers EMT but also significantly boosts their exosome secretion. This is consistent with previous studies showing that hypoxia modulates the secretome of cardiac progenitor cells by activating HIF-dependent pathways, thereby enriching their exosomal cargo with regenerative factors ^[20, 24]^. However, to our knowledge, this is the first study to specifically investigate the role of hypoxia-conditioned hEPs and to delineate the functional and molecular differences between normoxic and hypoxic exosomes.

Angiogenesis plays a pivotal role in the treatment of MI. Following an MI, the heart suffers ischemic damage, leading to impaired oxygen and nutrient supply to the affected myocardial tissue. Angiogenesis is crucial for restoring this blood supply, as it helps to revascularize the ischemic region, reducing tissue hypoxia and supporting cardiomyocyte survival ^[30]^. Our study found that miRNA-214-3p, enriched in Exo-H, can target VASH1 to promote angiogenesis in vivo and vitro. VASH1 is a protein that plays a critical role in regulating angiogenesis. It is primarily expressed in endothelial cells and has been identified as a potent negative regulator of angiogenesis. VASH1 inhibits angiogenesis by suppressing the activation of VEGF-mediated signaling pathways, which are crucial for endothelial cell proliferation, migration, and new blood vessel formation ^[31–33]^.

Mitochondria are the “power-houses” of cells, producing the energy necessary for a myriad of cellular processes ^[34]^. Accumulating experimental evidence has demonstrated that Drp1-mediated mitochondrial fission plays a crucial role in myocardial cell death during pressure overload and MI, ultimately leading to mitochondrial dysfunction and cardiomyocyte apoptosis ^[17, 35, 36]^. Inhibition of Drp1 using P110 (a peptide inhibitor that selectively blocks the Drp1–Fis1 interaction under pathological conditions) has been shown to prevent opening of the mitochondrial permeability transition pore, reduce infarct size, and ameliorate cardiovascular dysfunction in acute MI models. These findings suggest the potential clinical applicability of targeting Drp1 for therapeutic purposes ^[37]^. Our study reveals that miR-214-3p, which is enriched in Exo-H, can target MIEF2, thereby reducing Drp1 recruitment to mitochondria. This ultimately alleviates mitochondrial fission in cardiomyocytes following MI. The findings were further validated through TEM and mitotracker staining. Improvement of mitochondrial function inhibits mitochondrial outer membrane permeabilization (MOMP) and cytochrome c release by stabilizing mitochondrial membrane potential, enhancing ATP production, reducing ROS levels, maintaining calcium homeostasis, and upregulating anti-apoptotic Bcl-2 family proteins, thereby decreasing both caspase-dependent and caspase-independent apoptosis. So, in line with this, Exo-H and enriched miR-214-3p obviously reduced cardiomyocyte apoptosis under ischemic injury by targeting MIEF2 to alleviate mitochondrial fission and improve mitochondrial function.

MiR-214-3p has been shown to play a crucial role in various disease models ^[38–40]^. Concurrently, studies have demonstrated that exosomes derived from mesenchymal stem cells overexpressing miR-214-3p can effectively promote cardiac repair.^[41]^ In this study, we present a novel finding that miR-214-3p targets both VASH1 and MIEF2, thereby promoting angiogenesis and reducing mitochondrial fission in cardiomyocytes, which ultimately protects the heart following MI. This novel dual-targeting mechanism provides a profound insight into the specific therapeutic action of miR-214-3p in the treatment of MI, significantly broadening its potential clinical applications.

In conclusion, this study identifies hypoxia-conditioned hEP-Exo as a potent and multifaceted acellular therapy for ischemic myocardial injury. By leveraging the dual angiogenic and mitochondrial protective effects of miR-214-3p, Exo-H mitigates cardiomyocyte apoptosis, promotes vascular regeneration, and improves functional recovery after MI. These findings broaden the scope of regenerative cardiology therapy and pave the way for next-generation, cell-free, mechanistically defined exosome-based strategies. However, future work aimed at optimizing delivery, clarifying safety, and advancing clinical readiness will be essential to translate these insights into meaningful patient benefits.

## Data Availability Statement

The data that support the findings of this study are available from the corresponding author upon reasonable request.

## Funding

This work was supported by the National Key Research and Development Program of China (2022YFA1105100), the National Natural Sciences Foundation of China (82570339, 82270261, 82400333, and 82070260), the Eastern Talent Plan Leading Project in Shanghai (BJKJ2024014), the Academic Leaders Training Program of the Pudong Health Bureau of Shanghai (PWRd2024-02), the Natural Science Foundation of Shanghai Municipality (24ZR1459100), and the Peak Disciplines (Type IV) of Institutions of Higher Learning in Shanghai.

## Conflicts of Interest

NA

## Ethics approval statement

All animal procedures and protocols were performed in accordance with the Guide for the Care and Use of Laboratory Animals published by the U.S. National Institutes of Health and were approved by the Institutional Animal Care and Use Committee of Tongji University.

## Author Contributions

L.G. conceived and designed the research. Z.Q., Y.J., P.Z., H.L., Y.Y., and Y.G. contributed to the acquisition, analysis, and interpretation of the data. Z.Q., Y.J., and L.G. wrote and revised the manuscript. All authors read and approved the final manuscript.

---

**Figure S1.**
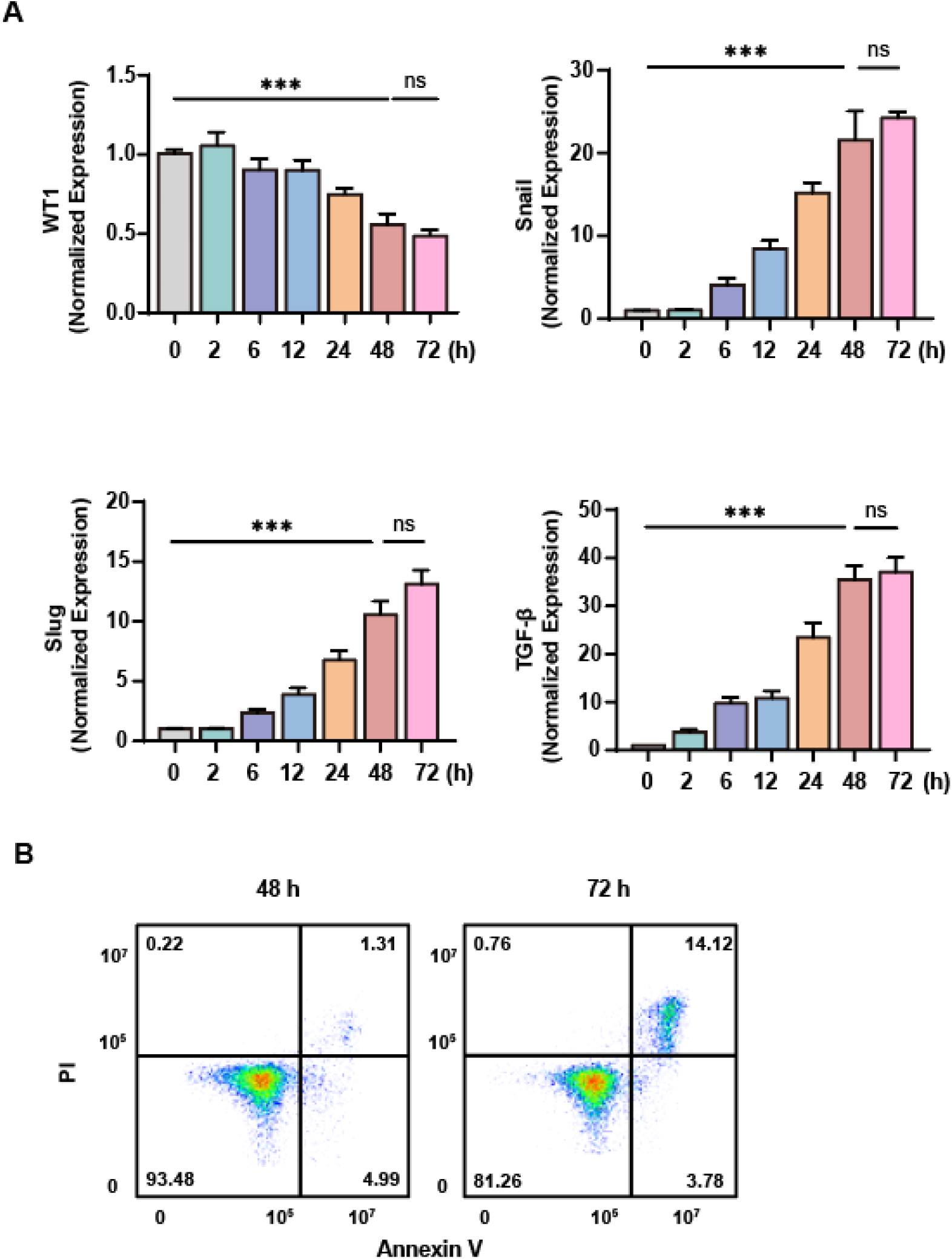
The status of hiPSC-derived epicardial cells (hEPs) at various time points. **(A)** Quantitative PCR (qPCR) was performed to analyze the expression of epithelial-mesenchymal transition (EMT)-related genes at various time points under hypoxic conditions. **(B)** Annexin V-PI flow cytometry was performed on hypoxic hEPs cultured for 48 and 72 h to assess cell apoptosis. Quantitative data are presented as mean ± SEM. For experiments involving three or more groups (A), a one-way ANOVA followed by Tukey’s post-hoc test for multiple comparisons was applied. \*\*\**P* < 0.001.

**Figure S2.**
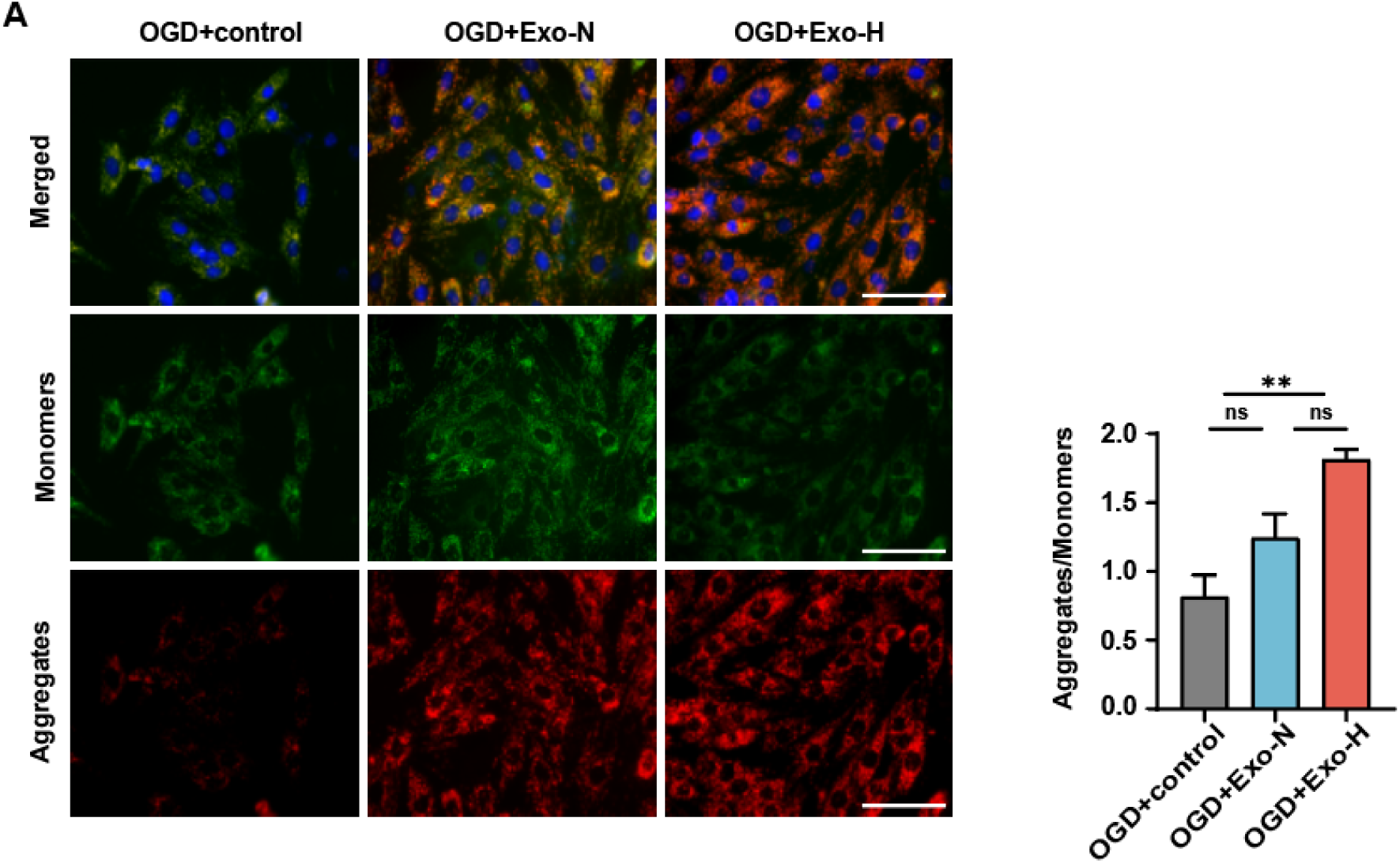
JC-1-based quantification of mitochondrial membrane potential in hiPSC-derived cardiomyocytes (hCMs). **(A)** The degree of mitochondrial depolarization was quantified by calculating the red-to-green fluorescence ratio. Quantitative data are presented as mean ± SEM. For experiments involving three or more groups (A), a one-way ANOVA followed by Tukey’s post-hoc test for multiple comparisons was applied. \*\**P* < 0.01, ns = not significant.

**Figure S3.**
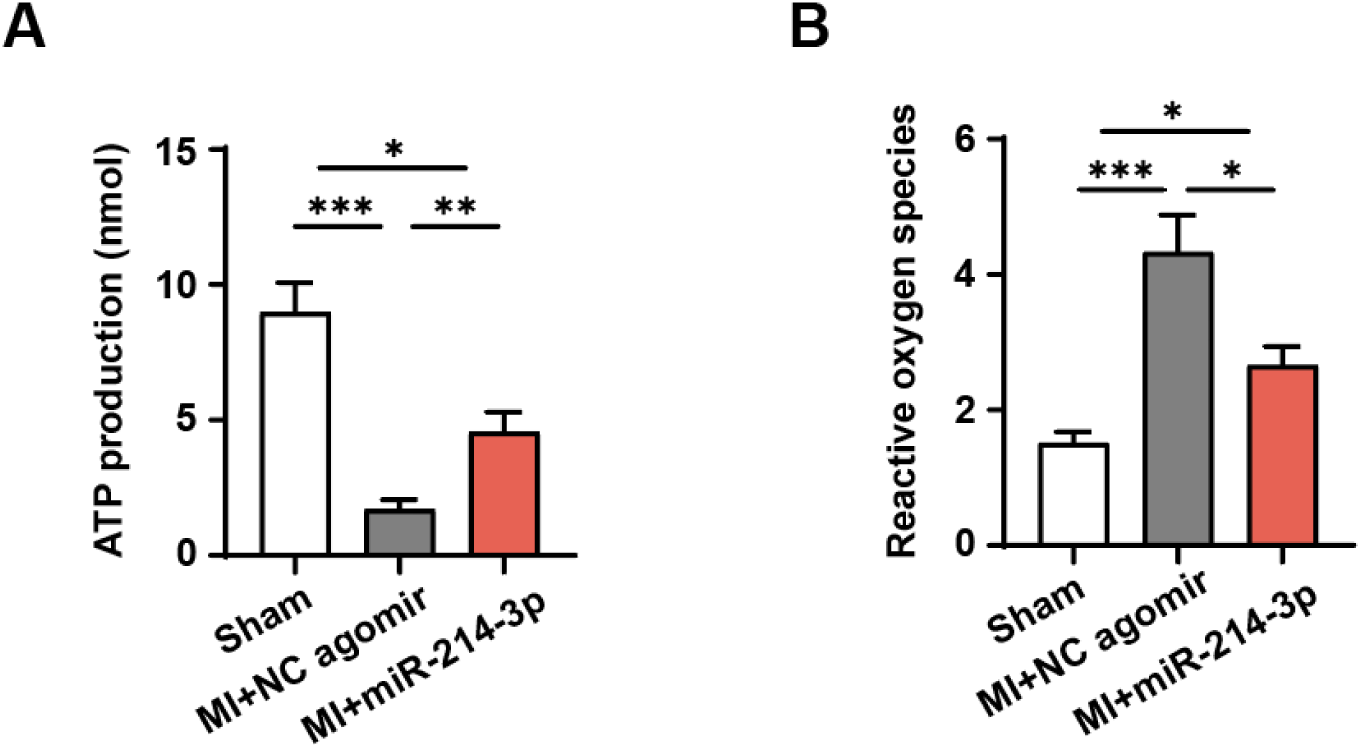
Mitochondrial dysfunction in cardiomyocytes post-MI. **(A)** ATP production in adult mouse cardiomyocytes (AMCMs) after MI. n=3. **(B)** ROS release in AMCMs after MI. n=3. Quantitative data are presented as mean ± SEM. For experiments involving three or more groups (A, B), a one-way ANOVA followed by Tukey’s post-hoc test for multiple comparisons was applied. \*\*\**P* < 0.001, \*\**P* < 0.01, and \**P* < 0.05.

## Notes

### Competing Interest Statement

The authors have declared no competing interest.

## Reference

1. Ziaeian, B. and G.C. Fonarow, Epidemiology and aetiology of heart failure. Nat Rev Cardiol, 2016. 13(6): p. 368–78.

2. Roger, V.L., Epidemiology of Heart Failure. Circulation Research, 2013. 113(6): p. 646–659.

3. Sanchez-Fernandez, C., et al., Regulation of Epicardial Cell Fate during Cardiac Development and Disease: An Overview. Int J Mol Sci, 2022. 23(6).

4. Wei, K., et al., Epicardial FSTL1 reconstitution regenerates the adult mammalian heart. Nature, 2015. 525(7570): p. 479-85.

5. Zhou, B., et al., Adult mouse epicardium modulates myocardial injury by secreting paracrine factors. J Clin Invest, 2011. 121(5): p. 1894–904.

6. Song, C., et al., Ammonium Persulfate-Loaded Carboxylic Gelatin-Methacrylate Nanoparticles Promote Cardiac Repair by Activating Epicardial Epithelial-Mesenchymal Transition via Autophagy and the mTOR Pathway. ACS Nano, 2023. 17(20): p. 20246–20261.

7. Luo, X.L., et al., hESC-Derived Epicardial Cells Promote Repair of Infarcted Hearts in Mouse and Swine. Adv Sci (Weinh), 2023: p. e2300470.

8. Sayed, A., et al., Hypoxia promotes a perinatal-like progenitor state in the adult murine epicardium. Sci Rep, 2022. 12(1): p. 9250.

9. Jing, X., et al., Hypoxia induced the differentiation of Tbx18-positive epicardial cells to CoSMCs. Sci Rep, 2016. 6: p. 30468.

10. Korvenlaita, N., et al., Dynamic release of neuronal extracellular vesicles containing miR-21a-5p is induced by hypoxia. J Extracell Vesicles, 2023. 12(1): p. e12297.

11. Bister, N., et al., Hypoxia and extracellular vesicles: A review on methods, vesicular cargo and functions. J Extracell Vesicles, 2020. 10(1): p. e12002.

12. Li, H., et al., Exosomes secreted by endothelial cells derived from human induced pluripotent stem cells improve recovery from myocardial infarction in mice. Stem Cell Res Ther, 2023. 14(1): p. 278.

13. Gao, L., et al., Therapeutic delivery of microRNA-125a-5p oligonucleotides improves recovery from myocardial ischemia/reperfusion injury in mice and swine. Theranostics, 2023. 13(2): p. 685–703.

14. Ma, T., et al., Therapeutic silencing of lncRNA RMST alleviates cardiac fibrosis and improves heart function after myocardial infarction in mice and swine. Theranostics, 2023. 13(11): p. 3826–3843.

15. Coller, H.A., MYC sets a tumour-stroma metabolic loop. Nat Cell Biol, 2018. 20(5): p. 506–507.

16. Wai, T. and T. Langer, Mitochondrial Dynamics and Metabolic Regulation. Trends Endocrinol Metab, 2016. 27(2): p. 105–117.

17. Ong, S.B., et al., Inhibiting mitochondrial fission protects the heart against ischemia/reperfusion injury. Circulation, 2010. 121(18): p. 2012–22.

18. Bao, X., et al., Directed differentiation and long-term maintenance of epicardial cells derived from human pluripotent stem cells under fully defined conditions. Nat Protoc, 2017. 12(9): p. 1890–1900.

19. 19. Bao, X., et al., Long-term self-renewing human epicardial cells generated from pluripotent stem cells under defined xeno-free conditions. Nat Biomed Eng, 2016. 1.

20. Wu, Q., et al., Extracellular vesicles from human embryonic stem cell-derived cardiovascular progenitor cells promote cardiac infarct healing through reducing cardiomyocyte death and promoting angiogenesis. Cell Death Dis, 2020. 11(5): p. 354.

21. Gao, L., et al., Large Cardiac Muscle Patches Engineered from Human Induced-Pluripotent Stem Cell-Derived Cardiac Cells Improve Recovery From Myocardial Infarction in Swine. Circulation, 2018. 137(16): p. 1712–1730.

22. Cao, N., et al., Highly efficient induction and long-term maintenance of multipotent cardiovascular progenitors from human pluripotent stem cells under defined conditions. Cell Res, 2013. 23(9): p. 1119–32.

23. Khan, M., et al., Embryonic stem cell-derived exosomes promote endogenous repair mechanisms and enhance cardiac function following myocardial infarction. Circ Res, 2015. 117(1): p. 52–64.

24. Gray, W.D., et al., Identification of therapeutic covariant microRNA clusters in hypoxia-treated cardiac progenitor cell exosomes using systems biology. Circ Res, 2015. 116(2): p. 255–63.

25. Bian, S., et al., Extracellular vesicles derived from human bone marrow mesenchymal stem cells promote angiogenesis in a rat myocardial infarction model. J Mol Med (Berl), 2014. 92(4): p. 387–97.

26. Nunnari, J. and A. Suomalainen, Mitochondria: in sickness and in health. Cell, 2012. 148(6): p. 1145–59.

27. Palmer, C.S., et al., MiD49 and MiD51, new components of the mitochondrial fission machinery. EMBO Rep, 2011. 12(6): p. 565–73.

28. Quijada, P., M.A. Trembley, and E.M. Small, The Role of the Epicardium During Heart Development and Repair. Circ Res, 2020. 126(3): p. 377–394.

29. von Gise, A. and W.T. Pu, Endocardial and epicardial epithelial to mesenchymal transitions in heart development and disease. Circ Res, 2012. 110(12): p. 1628–45.

30. Wu, X., et al., Angiogenesis after acute myocardial infarction. Cardiovasc Res, 2021. 117(5): p. 1257–1273.

31. Kobayashi, M., et al., Tubulin carboxypeptidase activity of vasohibin-1 inhibits angiogenesis by interfering with endocytosis and trafficking of pro-angiogenic factor receptors. Angiogenesis, 2021. 24(1): p. 159–176.

32. Sato, Y. and H. Sonoda, The vasohibin family: a negative regulatory system of angiogenesis genetically programmed in endothelial cells. Arterioscler Thromb Vasc Biol, 2007. 27(1): p. 37–41.

33. Chatterjee, S., Reversal of vasohibin-driven negative feedback loop of vascular endothelial growth factor/angiogenesis axis promises a novel antifibrotic therapeutic strategy for liver diseases. Hepatology, 2014. 60(2): p. 458–60.

34. Jin, J.Y., et al., Drp1-dependent mitochondrial fission in cardiovascular disease. Acta Pharmacol Sin, 2021. 42(5): p. 655–664.

35. Zhu, J., et al., MiR-181a protects the heart against myocardial infarction by regulating mitochondrial fission via targeting programmed cell death protein 4. Sci Rep, 2024. 14(1): p. 6638.

36. Zaja, I., et al., Cdk1, PKCdelta and calcineurin-mediated Drp1 pathway contributes to mitochondrial fission-induced cardiomyocyte death. Biochem Biophys Res Commun, 2014. 453(4): p. 710–21.

37. Disatnik, M.H., et al., Acute inhibition of excessive mitochondrial fission after myocardial infarction prevents long-term cardiac dysfunction. J Am Heart Assoc, 2013. 2(5): p. e000461.

38. Li, Y., et al., Downregulation of miR-214-3p attenuates mesangial hypercellularity by targeting PTEN-mediated JNK/c-Jun signaling in IgA nephropathy. Int J Biol Sci, 2021. 17(13): p. 3343–3355.

39. Dong, H., et al., miR-214-3p promotes the pathogenesis of Parkinson’s disease by inhibiting autophagy. Biomed Pharmacother, 2024. 171: p. 116123.

40. Fioravanti, A., S. Tenti, and S. Cheleschi, MiR-214-3p, a novel possible therapeutic target for the pathogenesis of osteoarthritis. EBioMedicine, 2021. 66: p. 103300.

41. Zhu, W., et al., Exosomes derived from mir-214-3p overexpressing mesenchymal stem cells promote myocardial repair. Biomater Res, 2023. 27(1): p. 77.

